# HIV-1 Envelope and MPER antibody structures in lipid assemblies

**DOI:** 10.1101/838912

**Authors:** Kimmo Rantalainen, Zachary T. Berndsen, Aleksandar Antanasijevic, Torben Schiffner, Xi Zhang, Wen-Hsin Lee, Jonathan L. Torres, Lei Zhang, Adriana Irimia, Jeffrey Copps, Kenneth Zhou, Young Do Kwon, William H. Law, Chaim A. Schramm, Raffaello Verardi, Shelly Krebs, Peter D. Kwong, Nicole A. Doria-Rose, Ian A. Wilson, Michael B. Zwick, John R. Yates, William R. Schief, Andrew B. Ward

## Abstract

Structural and functional studies of HIV Env as a transmembrane protein have long been complicated by challenges associated with inherent flexibility of the molecule and the membrane-embedded hydrophobic regions. Thus, most structural studies have utilized soluble forms where the regions C-terminal to the ectodomain are deleted. Here, we present approaches for incorporating full-length, wild-type HIV-1 Env, as well as C-terminally truncated and stabilized versions, into lipid assemblies, providing a modular platform for Env structural studies by single particle electron microscopy. We reconstituted a full-length Env clone into a nanodisc with MSP1D1 scaffold, complexed it with an MPER targeting antibody 10E8, and structurally defined the full quaternary epitope of 10E8 consisting of lipid, MPER and ectodomain contacts. By aligning this and other Env-MPER antibody complex reconstructions with the lipid bilayer, we observe evidence of Env tilting as part of the neutralization mechanism for MPER-targeting antibodies. We also adapted the platform toward vaccine design purposes by introducing stabilizing mutations that allow purification of unliganded Env with peptidisc scaffold.

## Introduction

HIV envelope glycoprotein (Env) is a homotrimeric transmembrane protein belonging to the class I viral fusion proteins. Binding of Env to host receptor CD4 and coreceptors CCR5 or CXCR4 leads to a cascade of conformational changes and eventually virus entry. Each of the three Env protomers are linked through a membrane proximal external region (MPER) to a single pass transmembrane domain (TMD) and an intracellular C-terminal domain (CTD), which play critical roles in fusion (Chen, 2019; Harrison, 2015; Santos da Silva et al., 2013). While several structures of isolated ectodomains, MPER peptides and TMDs have been determined using x-ray crystallography, cryo-EM and NMR, structural studies of the complete Env have been complicated by the challenging nature of the trimer (Chen, 2019; Ward and Wilson, 2017). Low expression levels and poor long-term stability of native Env together with structural flexibility and shedding of gp120 subunit (Hammonds et al., 2003), have led structural biologists to stabilize the trimer with mutations and/or antibodies so as to achieve higher resolution details. Structural intermediates have been captured after receptor binding and in complex with antibodies, which illustrate the intricately coordinated structural transitions of Env (Lu et al., 2019; Ozorowski et al., 2017; Tran et al., 2012). This flexibility is compounded when parts below the ectodomain are included, so structures of MPER, TMD and CTD have only been resolved in isolation using NMR (Chiliveri et al., 2018; Dev et al., 2016; Kwon et al., 2018a; Sun et al., 2008). In these studies, the MPER is found as a membrane-embedded amphipathic helix and trimeric protrusion from the membrane, TMD as a three-helix bundle or separate tilted helix, and the CTD as an elongated set of three amphipathic helices, leaving open questions as to how these conformations relate to the full Env trimer assembly. These structures have however provided important insights in the dynamic nature of these domains and show the capacity to adapt to different stages of the entry and fusion process akin to those observed in the ectodomain. Structures of PGT151 antibody stabilized ectodomains from C-terminally truncated JRFL, BG505 and B41 Env constructs and wild type, full length Env from PC64 and AMC011 donors confirmed the structural similarity to stabilized soluble Env, but due to structural flexibility in micelle-embedded domains, the MPER, TMD and CTD have remained unresolved (Cao et al., 2018; Lee et al., 2016; Rantalainen et al., 2018; Torrents de la Pena et al., 2019).

A multitude of broadly neutralizing antibodies (bNAbs) have now been characterized, targeting various sites on HIV-1 Env (Haynes and Mascola, 2017; Sok and Burton, 2018). Structures of these bNAbs in complex with Env have paved the way for structure-based vaccine design by facilitating the identification of stabilizing mutations that allow large scale expression and purification of the unliganded ectodomain (Dey et al., 2018; Guenaga et al., 2017; Joyce et al., 2017; Klein et al., 2013; Kulp et al., 2017; Kwon et al., 2015; Ringe et al., 2017; Rutten et al., 2018; Schoofs et al., 2019; Sliepen et al., 2019; Torrents de la Pena and Sanders, 2018; Ward and Wilson, 2017; Yuan et al., 2019). The epitopes of MPER-targeting bNAbs were however truncated along with TMD and CTD from almost all of these Envs to increase solubility and stability and, therefore, remains perhaps the least understood epitope. Well-known members of the MPER bNAb family are antibodies 10E8 and 4E10, which share common features and high neutralization breadth over different HIV strains (Cardoso et al., 2005; Huang et al., 2012; Stiegler et al., 2001). More recently, a new MPER targeting antibody lineage PGZL1 from donor PG13, and three lineages from donor RV-217 (VRC42, VRC43, VRC46) with outstanding breadth were discovered. These studies describe in more detail the maturation of MPER-targeting antibodies (Krebs et al., 2019), [Zhang, L. et. al. An MPER Antibody Neutralizes HIV-1 Using Germline Features Shared Among Donors. Nat Commun. 2019. In Press]. While the mature PGZL1 neutralized 84% of the 130-virus panel with ~21% heavy chain (HC) and ~13% light chain (LC) somatic hyper mutation (SHM) at nucleotides level, the recombinant sub-lineage variant H4K3 (17% HC, 12% LC SHM) was able to neutralize 100% of the panel and importantly, the germline reverted variant neutralized 12% of the panel. In RV-217, all three lineages matured with lower SHM (9-13%) with up to 96% neutralization breadth, whereas the VRC42 lineage reached 50% neutralization breadth with only 2% SHM. These impressive bNAbs together with the high conservation of MPER sequence has stimulated renewed interest in MPER-targeting vaccine design and the use of MPER antibodies for post-exposure prophylaxis. Crystal structures of many MPER Fabs have been solved, alone and in complex with MPER peptide and/or with additional short-tailed lipid headgroups (Irimia et al., 2016; Irimia et al., 2017; Krebs et al., 2019; Williams et al., 2017), [Zhang, L. et. al. An MPER Antibody Neutralizes HIV-1 Using Germline Features Shared Among Donors. Nat Commun. 2019. In Press], providing valuable details of MPER peptide and membrane lipid recognition. For example, in the case of 10E8, residues in CDRL1 and CDRH3 were shown to bind to phosphatidic acid and phosphatidylglycerol headgroups. Two intermediate resolution cryo-EM studies of MPER-targeting antibodies (10E8 and PGZL1) in the context of the trimeric ectodomain provide insights into antibody approach angle, steric obstruction by glycans and antibody induced lifting of Env from lipid surface (Lee et al., 2016), [Zhang, L. et. al. An MPER Antibody Neutralizes HIV-1 Using Germline Features Shared Among Donors. Nat Commun. 2019. In Press]. Despite efforts to better understand the MPER antibodies, the full quaternary epitope has remained elusive, emphasizing the need to study structures of these antibodies in more native environments.

Reconstitution of membrane proteins into lipid nanodiscs in combination with recent advances in cryo-EM has shown great promise as a tool for structural biology of membrane proteins (Denisov and Sligar, 2016; Efremov et al., 2017). This method was made possible by the introduction of apolipoprotein A based scaffold proteins and associated nanodisc assembly methodology in the early 2000’s (Bayburt et al., 2002). Since then several other scaffold types have been introduced, all of which facilitate spontaneous assembly of the target protein into lipid bilayer discs upon detergent removal. In addition to rendering hydrophobic and transmembrane molecules to be essentially as manageable as soluble proteins, the nanodisc technology offers exceptional versatility for experimental design, allowing different disc diameters and lipid compositions to be co-assembled with the target molecule.

In this work, we present approaches to study HIV Env in membranous environments by assembling two full length (FL), wild type Envs from PC64 (clade A) and AMC011 (clade B) donors and a C-terminally truncated BG505 Env (clade A) into detergent-lipid micelles, bicelles, nanodiscs and peptidiscs. In combination with single particle EM analysis, these protein-lipid assemblies provide a tool for studying HIV Env in a lipid bilayer as well as the binding mechanism of MPER bNAbs. In the lipid bilayer systems, MPER bNAbs induce tilting of Env relative to the membrane surface, forming a wedge between the ectodomain and the lipid surface before lifting the Env off the membrane. In addition, by complexing the AMC011FL nanodisc with 10E8 Fab, we were able to characterize the full tripartite quaternary epitope consisting of lipid, peptide and glycan contacts. Finally, we show that this methodology can be adapted for vaccine engineering by introducing stabilizing mutations into a C-terminally truncated BG505 construct, allowing presentation of the full array of bNAb epitopes, including MPER, without the need for a stabilizing antibody.

## Results

### Env incorporation into detergent-lipid micelles, bicelles and nanodiscs

Different assembly pathways were experimentally assessed to establish a modular platform for studying Env and the neutralization mechanism of MPER antibodies in lipid environments (Figure 1, Figure S1). To build upon the detergent-lipid micelle approach described earlier (Blattner et al., 2014; Lee et al., 2016; Rantalainen et al., 2018; Torrents de la Pena et al., 2019), through a more complete detergent removal by and addition of scaffold proteins MSP1D1 or peptidisc, Env could be incorporated into bicelles and nanodiscs with various lipid compositions (Figure 2A). In detergent-lipid micelle approach lipid molecules are exchanged by partial detergent removal, leading to unstable (~1-2 days) complexes but suitable for high resolution determination of the ectodomain by cryo-EM. In nanodiscs, a complete detergent removal is done over ~48 hours in presence of MSP1D1 scaffold protein, leading to a stable lipid bilayer encircled by the scaffold. The peptidisc approach follows same principles but scaffold is now a short bi-helical, engineered peptide (Carlson et al., 2018). In the lipid bicelle approach, similar detergent removal leads to heterogeneously sized bicelles capped by lipid molecules and/or scaffold. Formation of bicelles versus nanodiscs was concluded to be largely dependent on the Env clone. Full-length PC64 Env (PC64FL) Env preferred the formation of larger bicelle assemblies with multiple Envs compared to AMC011FL, BG505∆CT and BG505-ST-710 (Figure 1B), which predominantly resulted in smaller discs with one or two Envs. Selection of lipids also affected the size distribution of the assemblies, although we could not systematically define the effect of lipid composition (Figure S1A). Env occupancy varied from one to two in nanodiscs to between two and four in larger bicelles. Size exclusion chromatography did not efficiently separate the different species and, in most samples, nanodiscs and bicelles were pooled. Different Env occupancies could, however, be easily separated computationally during 2D and 3D classification in EM data processing. Nanodiscs had an expected diameter of 10 nm defined by the MSP1D1 scaffold, whereas bicelle size varied from ~ 15 to ~ 25 nm (Figure 2B and D). Incorporation of lipid molecules was confirmed with mass-spectrometry, where five out of seven added lipid types could be confirmed (Figure S1C, 1D and 1E). The Env incorporation and recovery ratio varied between 5-50% and was reproducibly higher with the peptidisc scaffold. Incorporation was also noted to be more efficient and reproducible in smaller, 50-100µL reaction volumes. In control reactions without scaffold protein, less stable proteoliposomes were formed (Figure S1B). In the absence of lipids and scaffold, rosettes of two or more Envs were formed leading to aggregation after prolonged incubation (~1-2 days, Figure S1B). PC64FL and AMC011FL Env nanodiscs and bicelles remained intact at +4°C for up to 4 months but could not be recovered after freezing or complete dehydration (Figure S1B). Samples were primarily assessed using negative-stain and cryo-EM single particle image analysis (Figure S2).

**Figure 1.**
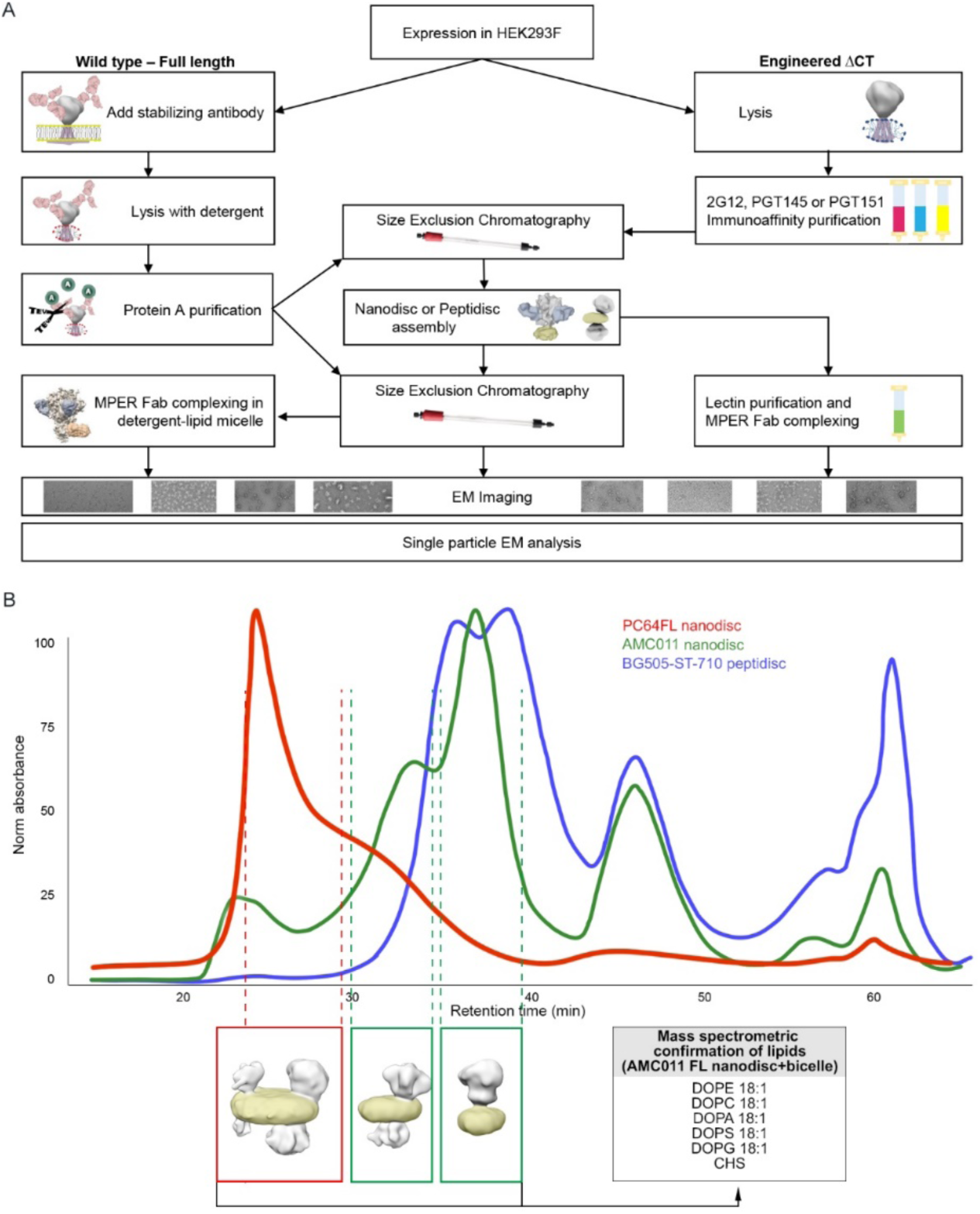
Preparation of Env lipid assemblies. A) Overview of the workflow used to generate different lipid assemblies. B) Typical size exclusion chromatograms of different Env assemblies and 3D reconstructions below the chromatogram peaks illustrate the corresponding forms of the assembly with the lipid bilayer highlighted in yellow. Incorporation of lipids was confirmed by mass spectrometry from a pool of AMC011 FL nanodiscs and bicelles.

**Figure 2.**
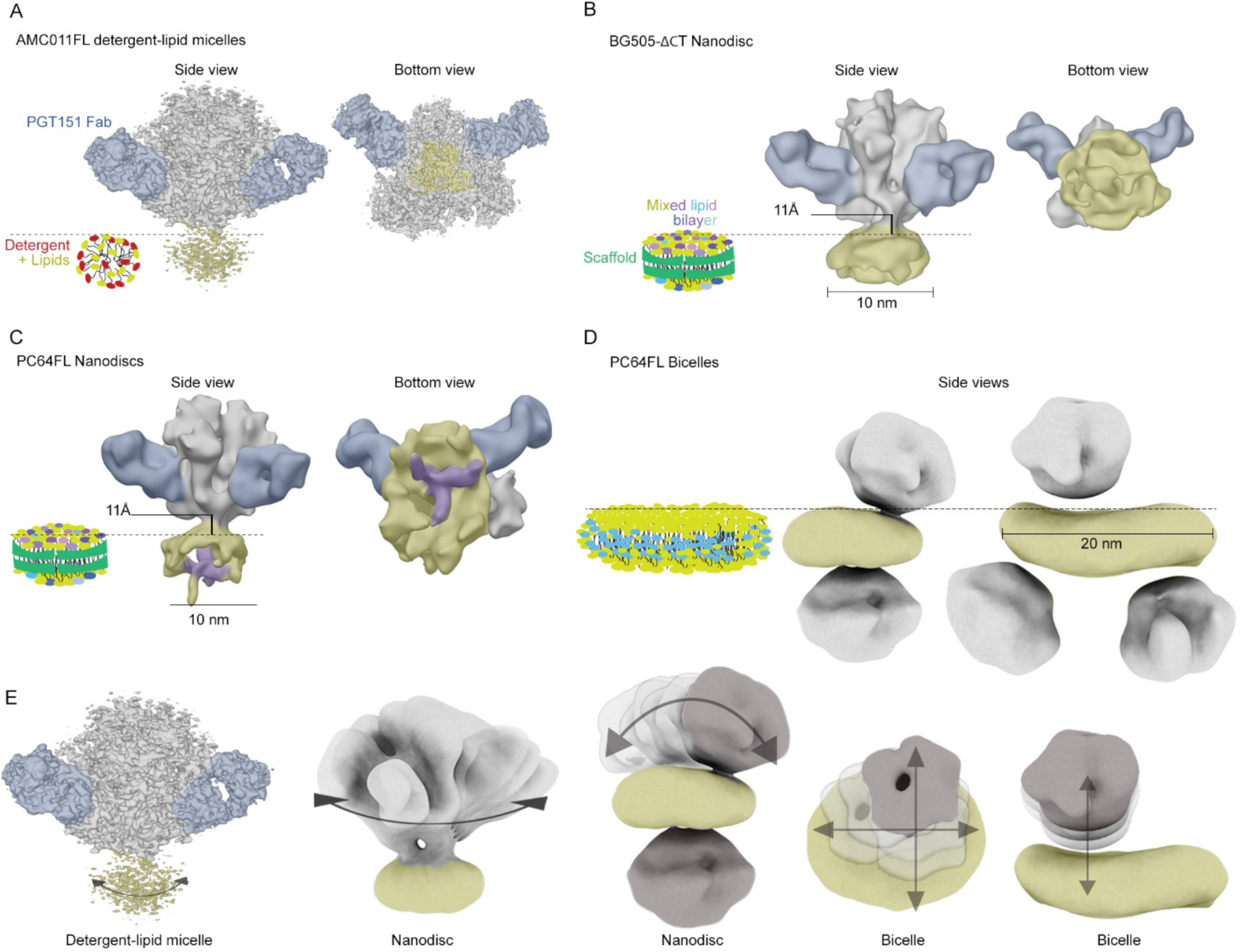
Comparison of different sample preparation approaches. Stabilizing PGT151 Fab highlighted in blue. Different assembly types are presented with cartoons next to negative stain and cryo-EM 3D reconstructions. Env ectodomain is highlighted in gray, PGT151 in blue, micelle and lipid bilayer in yellow. A) The ectodomain can be reconstructed to 3 – 5Å resolution from all lipid-detergent micelle samples. B and C) Cryo-EM reconstructions of nanodiscs with height of the ectodomain from the bilayer surface indicated and density that passes through the disc of PC64FL is highlighted in purple. D) In bicelles, additional compositional heterogeneity limits the analysis to negative-stain EM reconstructions. E) Summary of the rotational, positional and compositional heterogeneity.

### Env displays several degrees of flexibility in lipid assemblies

In all detergent-lipid micelle PC64 and AMC011 FL samples, the PGT151-stabilized ectodomain could be reconstructed to 4-5 Å resolution (Figure 2A). In the nanodisc sample of BG505∆CT, the PGT151-stabilized ectodomain was reconstructed to 4.6 Å (Figure S2A), but the global resolution ranged between 9-12 Å when disc and regions below the ectodomain were included (Figure 2B and C, table S1). Nonetheless, this resolution enabled positioning of the bilayer surface and measurement of stable part of ectodomain height from the membrane surface (ending at Asp664). In both BG505∆CT and PC64FL, the ectodomain was 11 Å from the bilayer surface (Figure 2B and C). The PC64FL nanodisc reconstruction at 9 Å resolution contained a continuous density emanating from the bottom of the ectodomain and spanning the bilayer (Figure 2C, Figure S3B). In bicelles, two to four Envs were incorporated thereby greatly increasing sample heterogeneity and limiting the studies to negative-stain EM analysis (Figure 1B, 2D, S2A, S5). In some 2D and 3D classes, some bicelle curvature was also observed. In summary, Env showed several degrees of rotational and positional flexibility as well as compositional heterogeneity in nanodiscs and bicelles, limiting high resolution structure determination but likely reflecting the native flexibility of the trimer on the surface of membranes (Figure 2E).

### Env-MPER Fab complexes in detergent-lipid micelles show heterogenous Fab positioning and TMDs crossing at residue R696

We next attempted to stabilize the flexible parts of Env and improve the epitope definition by addition of different MPER-targeting antibody Fabs in mixed detergent-lipid micelles (Figure 3). In all tested combinations of FL Env and MPER Fab, cryo-EM analysis showed no stable, high-resolution structural features for MPER, TMD or CTD similarly to earlier cryo-EM studies (Figure 3A) (Lee et al., 2016; Rantalainen et al., 2018; Torrents de la Pena et al., 2019). One of the tested antibodies involved a variant of 10E8, called 10E8v4_5R+100cF, which was designed to have improved membrane-interaction capacity and which showed 10-fold high neutralization potency (Kwon et al., 2018b). This antibody also failed to show stable, high-resolution structural features suggesting that stable lipid headgroup contacts are not fully recovered in this approach. Despite collecting large cryo-EM datasets of over 1 million. particles, the liganded complexes could not be refined beyond 6-9 Å resolution for the regions below the ectodomain due to structural and compositional heterogeneity. The flexibility was further emphasized with multibody refinement of AMC011FL in complex with one copy of PGZL1 Fab (Video S1), where Fab and micelle show both horizontal and vertical movement in relation to the ectodomain. Similar evidence was provided by superimposition of different Fab occupancy classes from AMC011FL-VRC42.01 dataset showing a continuous Fab position around the micelle (Figure S4A). By collecting larger dataset and increasing the particle numbers of PC64FL in complex with VRC42.01 Fab, a subset of particles could be classified that contained continuous density from HR2 to MPER and into the TMD (Figure 3B, Table S1). Although the resolution did not enable *de novo* building of an atomic model, fitting in high-resolution structures of the complex components was sufficient to model the MPER and TMD. Interestingly, the TM helices crossed the micelle in a tilted fashion forming an X shape with TMDs from adjacent protomers crossing at approx. 75 ° angle (Figure 3B). The helices crossed in the micelle at the conserved R696, a residue previously shown to be important for modulating conformational changes of the TMD (Cooper et al., 2018; Hollingsworth et al., 2018; Wang et al., 2019).

**Figure 3.**
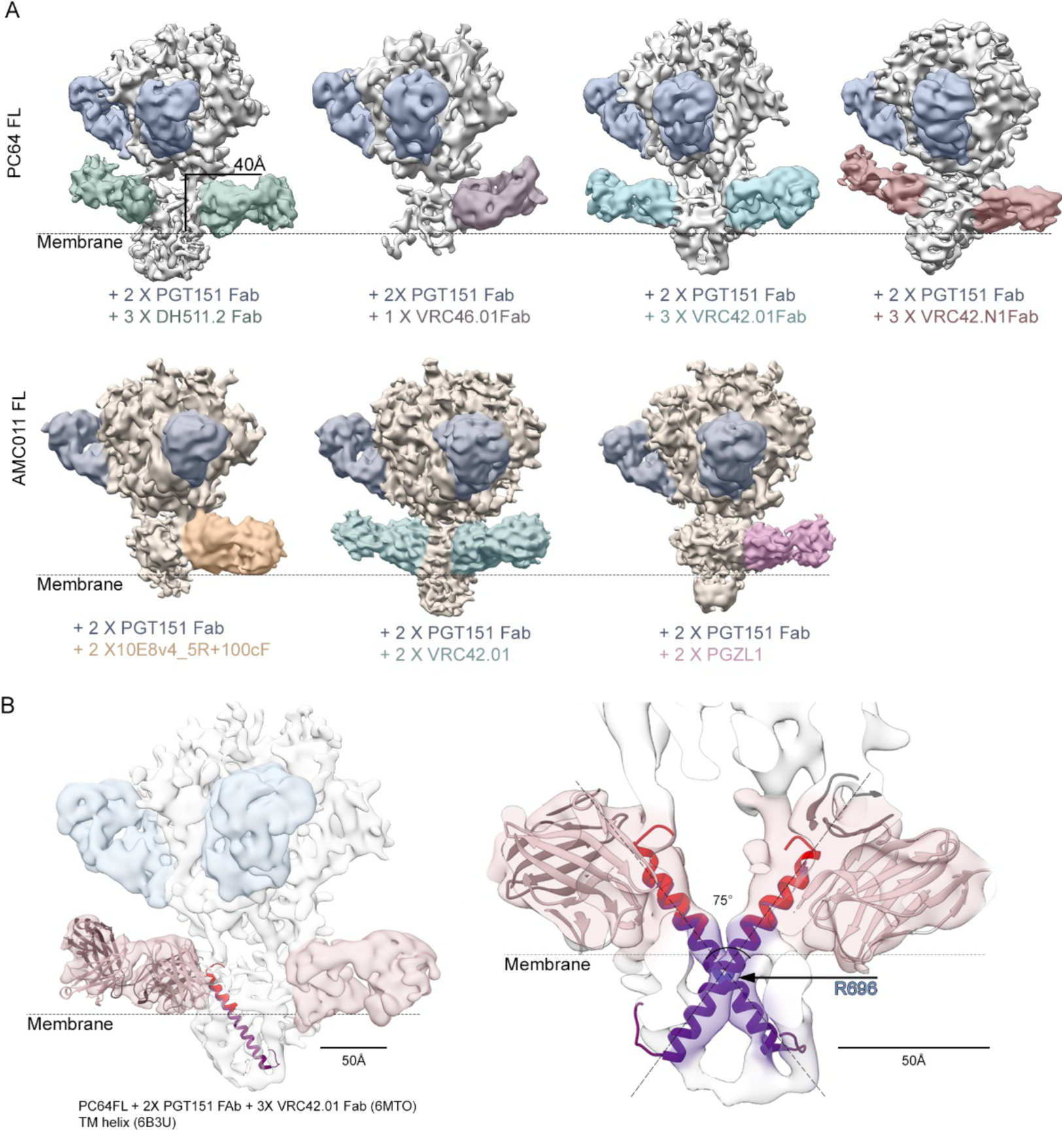
Cryo-EM reconstructions of FL Env-MPER Fab complexes in detergent-lipid micelles. A) PC64 and AMC011FL Envs with a panel in of MPER targeting Fabs showing variable Fab positions and occupancies. The estimated membrane position is indicated as well as the height of the structurally stable part of ectodomain (ending at Asp664) from the membrane surface. Particle classes with one, two and three Fabs could be classified from most of the datasets with all showing similar flexibility and heterogenous density in the micelle. When second or third MPER Fab is not visible, it is bound behind the micelle, pointing away from the viewer. B) In the complex between PC64FL and VRC42.01 Fab, the MPER density could be followed through to the TMD as continuous density allowing docking of crystal structure of the Fab (PDB: 6MTO) and NMR structure of tilted TMD helices (PDB: 6B3U). The position of R696 as the crossing point of the helices is indicated.

### EM analysis of MPER antibodies bound to bicelle and nanodisc incorporated Env reveals a wedging and tilting component in the binding mechanism

We next analyzed the MPER antibody binding mode in lipid bilayer assemblies with varied lipid content. In all tested combinations, addition of the Fab to Env embedded in a lipid bilayer (nanodisc or bicelle) led to displacement of Env to the side of the assembly and tilting of Env in relation to bilayer (Figure 4, Figure S4C) and Figure S5). This suggested that Env tilting is a common phenomenon of MPER antibody binding and prompted us to align the complexes to the bilayer instead of the ectodomain. By comparing the bilayer assemblies to detergent-lipid reconstructions, we also noted that the angle between the ectodomain and the Fab is identical between the two assembly types (Figure 4B). To confirm the tilting component of the MPER binding and to further elucidate the tripartite quaternary epitope of the antibody, we then assembled AMC011FL in nanodiscs for cryo-EM analysis with a lipid mixture roughly following the lipid composition earlier determined for HIV particles (Lorizate et al., 2013). We chose to complex the nanodisc with 10E8 Fab as this is the most characterized MPER-targeting antibody. The complex was frozen on graphene oxide grids to improve particle orientation distribution for cryo-EM analysis. These improvements allowed collection of data with low concentration of sample (~0.1mg/mL) and an adequate number of particles to classify stable particle subsets with different Fab occupancies from relatively small dataset (128,594 particles, Figure 4C, Figure S4B). Due to steric constraints introduced by the lipid bilayer, 10E8 now showed a fixed position in comparison to detergent-lipid approach, which in turn improved the 3D classification accuracy of the particles (Figure S4B). When the reconstructions were aligned to a simulated bilayer model, the Env ectodomain with one and two 10E8 Fabs was ~25 Å away and tilted 110° in relation to the bilayer surface, confirming the tilting observed with bicelles and negative-stain EM data (Figure 4D), whereas the third Fab led to an ~10° more vertical Env orientation. The height of the stable part of Env ectodomain from bilayer surface was now 30 Å compared to ~18 Å with one or two Fabs or 11Å in PC64FL and BG505∆CT nanodisc without MPER Fabs. The highest global resolution (~5 Å) was obtained with the complex containing one copy of PGT151 Fab and three copies of 10E8 Fab, which also had the largest number of particles (Figure 4E). By fitting the crystal structure of 10E8 together with Env protomers in the reconstruction, we generated a hybrid model allowing the definition of the tripartite quaternary epitope and an estimate of the contacting residues (Figure 4E and table 1, PDB ID: xxxx). While local resolution estimations showed stabilization of the epitope between Fab A and Env up to 5 Å resolution, the disc, TMD and CTD were less homogenous (Figure 4F). The TMDs could, however be resolved as a continuous density so that the Fab A and B complex had vertical TMD density and Fab C a tilted TMD, similar to micelle-embedded TMD in the PC64FL-VRC42.01 Fab complex (Figure 4G, Figure 3B). AMC011FL nanodisc bound to 10E8 had two distinct binding modes with different binding angles and ectodomain contacts. Fab A and B had a more vertical orientation in relation to the ectodomain and the third (Fab C) tilted orientation was influenced by the proximal PGT151 (Figure 4H). Based on fitting of ectodomain and Fab structures in the map, contacts to the ectodomain α8-helix (gp41) and gp120 by CDRH1 and heavy chain FR3 remained further apart in Fab C compared to A and B (Figure 4E). As PGT151 does not affect the orientation of 10E8 directly by steric blocking, we assume the differing orientation of Fab C is rather mediated by the stabilization or asymmetric distortion of the protomer-protomer interface induce by PGT151. When fitted positions of 10E8 Fab-MPER peptide crystal structures in AMC011 nanodisc were compared to ones in the JRFL∆CT micelle complex (Lee et al., 2016), the membrane surface facing side of the MPER peptides were approximately 10Å closer to each other (4I).

**Table 1.**
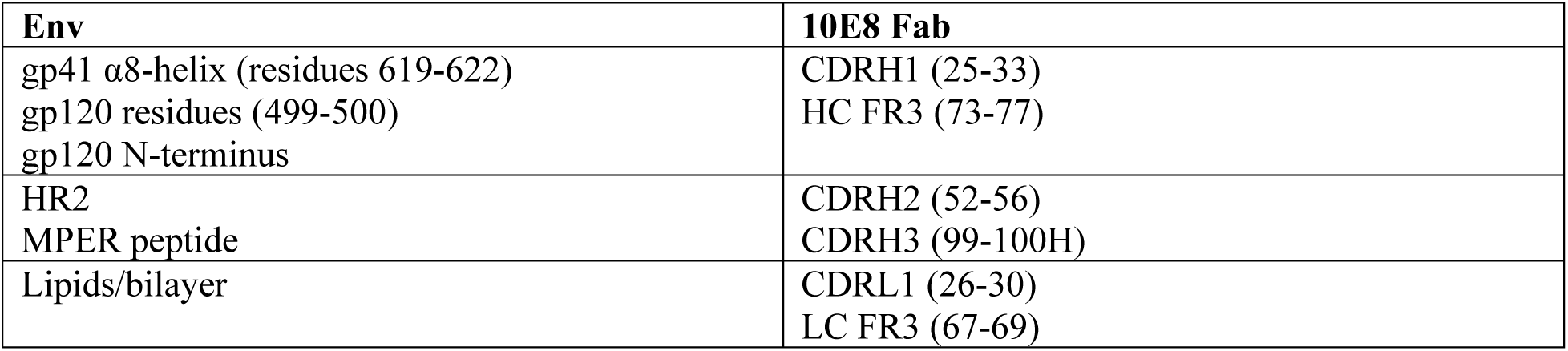
Contacting regions between Env and MPER-targeting antibody 10E8 mapped from the cryo-EM reconstruction of AMC011FL nanodisc in complex with the Fab.

**Figure 4.**
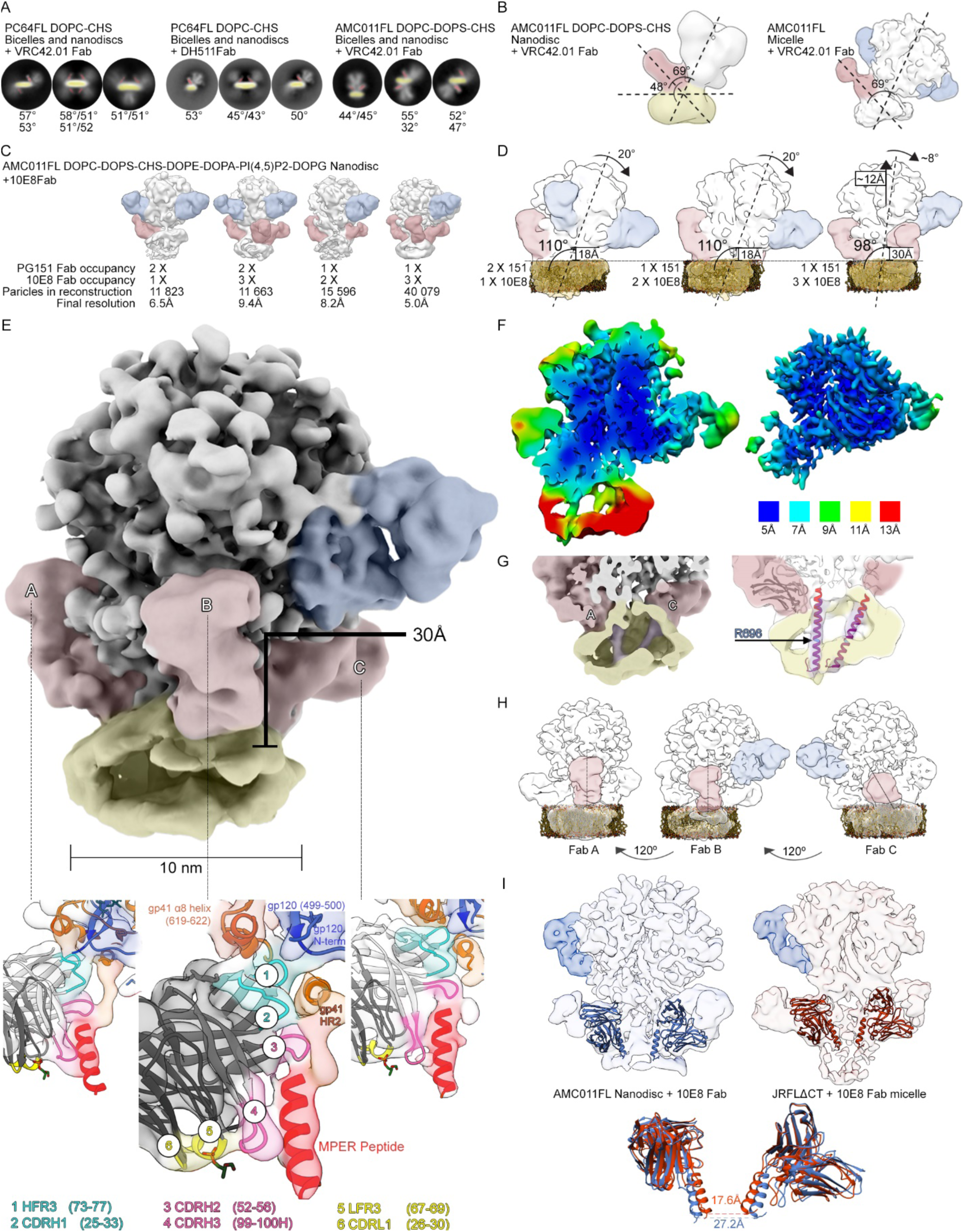
Analysis of FL Env in lipid bicelles and nanodiscs in complex with MPER targeting Fabs. A) Representative 2D class averages from negative-stain EM data from lipid assemblies of PC64FL and AMC011FL in complex with MPER-targeting antibodies showing tilting of ectodomain and estimated Fab angles in relation to bilayer. Lipid bilayer is highlighted in yellow and MPER Fab in red. B) Comparison of VRC42.01 Fab binding angle in nanodisc and micelle. PGT151 Fab is highlighted in blue throughout the figure. C) Low-pass filtered cryo-EM reconstructions of AMC011FL nanodisc in complex with 10E8 Fab with different Fab occupancies. Highest resolution and particle count were obtained with complex containing one copy of PGT151 Fab and three copies of 10E8 Fab, which is used for panels D through I. D) Low-pass filtered reconstructions with one, two or three copies of 10E8 Fab showing first tilting and then lifting of Env with addition of third 10E8 Fab. Approximate tilt degree of Env in relation to bilayer surface is indicated as well as change from vertical position. E) Highest resolution reconstruction and epitopes of the three Fabs with 10E8 Fab crystal structure docked in (PDB ID:5T80). Epitope components are highlighted as indicated in the panel below. Distance from bilayer surface is also indicated. F) Local resolution estimation showing up to 5Å resolution in the ectodomain and stabilized 10E8 Fab epitope. G) Horizontal (Fab A) and tilted (Fab C) Fab stabilized TMD orientation are highlighted in purple. H) Two Fab orientations and dependency of Fab C on the PGT151 position is highlighted in red in low-pass filtered maps. I) Comparison of 10E8 Fab docking in AMC011FL nanodisc and JRFL∆CT micelle reconstructions (EMD-3312) showing a ~10 Å change in the distance between the MPER peptides (residue Gln135 in PDB 5T80).

### Method adaptation to immunogen design

To overcome the requirement of using antibodies as a purification and stabilization reagent, and to adapt our methods for immunogen design purposes, we introduced stabilizing mutations to Env. BG505 Env was engineered with SOSIPv5.2 (Torrents de la Pena et al., 2017) and MD39 mutations (Steichen et al., 2016) that have previously been demonstrated to increase the stability and expression of soluble trimeric ectodomain of BG505. In addition, the construct was truncated at the C-terminus at residue 710 to eliminate unwanted epitopes. This construct was named BG505-ST-710. Soluble version of this construct was generated by TMD deletion and named BG505-ST-664. The introduced modifications allowed purification workflows similar to soluble SOSIP versions and production of unliganded TMD containing trimer. To reduce the introduction of potentially immunogenically active elements from the scaffold protein and to improve assembly efficiency, we utilized apolipoprotein A1 derived bi-helical peptidisc scaffold (Carlson et al., 2018). Assemblies with this peptidisc scaffold showed higher assembly efficiency with similar Env occupancy per disc and MPER binding modes as with PC64FL and AMC011FL nanodiscs apart from MPER Fab complex classes where Env was tilted over the side of the disc, most likely resulting from the lack of anchoring by CTD (Figure 5A, Figure S5). Binding of this construct to a panel of antibodies was tested by bio-layer interferometry (BLI, Octet), using lectin-based Env capture (Figure 5B). Quaternary-specific Fab PGT145, which preferentially binds to correctly folded trimers, bound to Env peptidiscs with similar binding levels (Figure. 5B) and affinity (Figure. 5C) as to soluble BG505-ST-664. Binding levels of PGT151 to Env peptidiscs were lower compared to the soluble trimer, but the affinities were similar (Figure. 5C). Non-neutralizing antibody B6, which binds only to non-native forms of Env, and 39F, which recognizes non-neutralizing epitopes in the V3 loop, showed minimal reactivity to both samples (Figure. 5B). 10E8 bound only to Env peptidiscs, but not to BG505-ST-664, which lacks the MPER epitope (Figure. 5B and D). Taken together, these data confirm that Env-peptidiscs are predominantly in native pre-fusion conformations similar to soluble BG505-ST-664.

**Figure 5.**
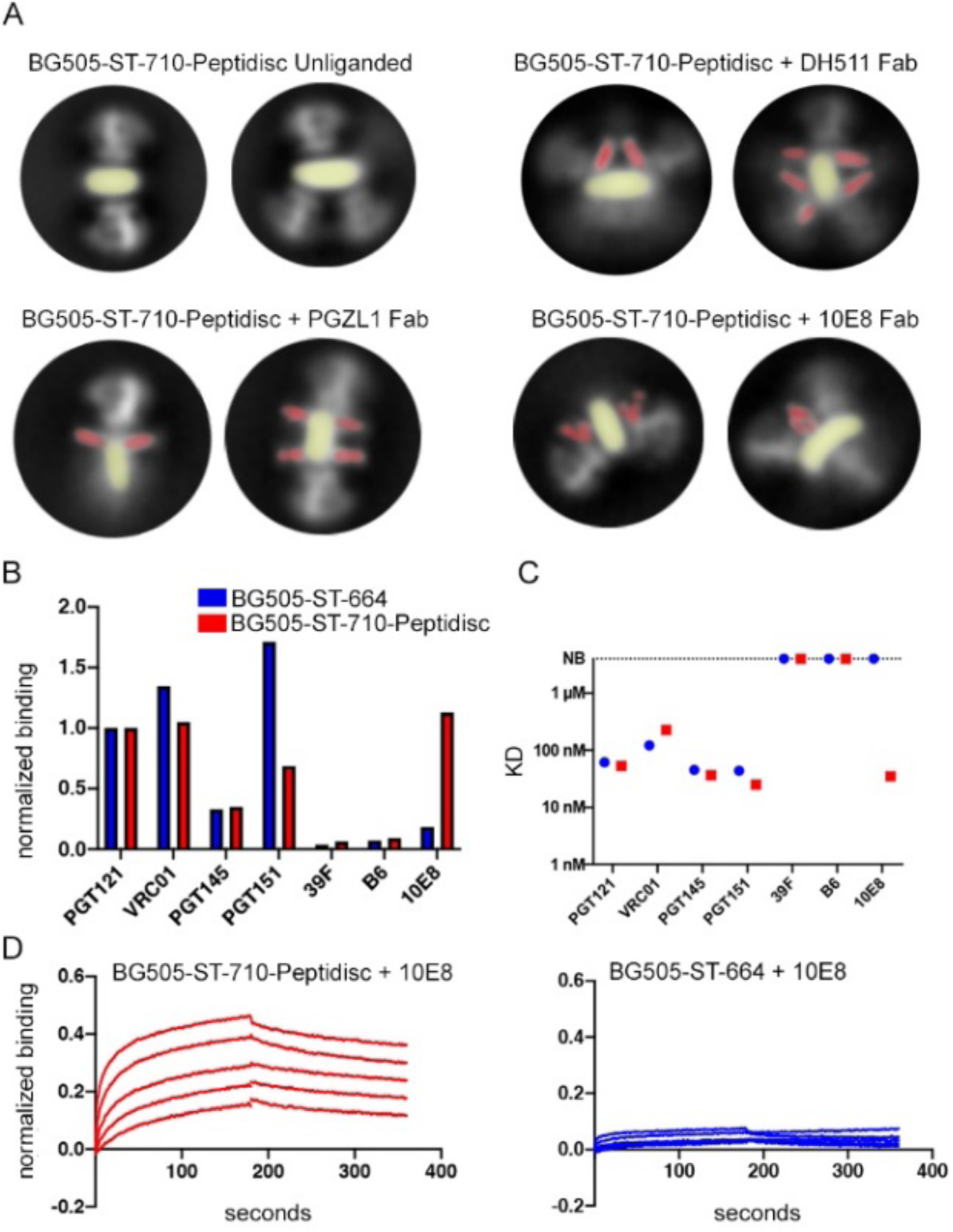
Assembly of vaccine design adapted BG505-ST-710 into peptidiscs. A) Examples BG505-ST-710 peptidiscs with and without the addition of MPER-targeting Fabs as determined from negative-stain EM 2D class averages. B-D) Antigenic profiling of BG505-ST-710 peptidiscs compared to equivalent soluble Env (BG505-ST-664) by lectin capture-based bio-layer interferometry. B) Maximum binding signals of the indicated Fabs are shown after normalization to the respective PGT121 signal. C) Kinetic analysis of all Fabs tested. NB: no binding D Exemplary titration series of MPER-specific Fab 10E8 against BG505-ST-710-Peptidisc and corresponding BG505-ST-664 construct.

## Discussion

Here we developed lipid assembly methods and demonstrate that these, in combination with single particle EM analysis, are well suited to improve our understanding of the structural dynamics of the HIV-1 Env glycoprotein. The system is currently limited to either PGT151 stabilized FL, wild type constructs or constructs with stabilizing mutations in the ectodomain. In addition, the assemblies are small patches of membrane that may limit the mobility and flexibility of the Env trimer. Nevertheless, the system is a step toward more native environment and open new possibilities for studying Env structure in lipid bilayers by single particle EM analysis. In addition to controlled lipid composition and the testing new MPER antibodies such as 10E8v4_5R+100cF with improved lipid contacts (Kwon et al., 2018b), these membrane embedded Envs may allow structural studies of the effect that CD4 receptor engagement has on the neutralization mechanism of the MPER bNAbs, which thus far has not been possible. As these complexes are likely heterogenous, excellent scalability and the classification power of single particle EM analysis becomes an important factor in addition to being able to yield high resolution structural data. Systematic studies of lipid composition, larger macromolecular complexes, as well as immunization trials, will therefore be the subject of future research.

Our studies demonstrate that the ectodomain of membrane embedded Envs are stable when bound to PGT151 or when stabilized with mutations, while the MPER and TMD exhibit a large degree of positional and local structural heterogeneity reflective of local dynamics near the membrane. Structural heterogeneity could be reduced by addition of MPER Fabs, or by incorporation into a nanodisc. Although we did not achieve high enough resolution to build atomic models for MPER, TMD or CTD, we could extract new details of these typically dynamic domains through 3D classification and by docking high resolution structures of complex components into the reconstructions.

For the PC64FL micelle in complex with VRC42.01 Fab and for the AMC011FL nanodisc with 10E8 Fab, densities likely corresponding to the MPER -bound TMD could be traced through the micelle and nanodisc respectively. Interestingly, a tilted orientation of TMDs could be determined in micelles with a crossing point at conserved residue R696. Influenza hemagglutinin was shown to have similar TMD dynamics with tilted and straight orientations, suggesting that these could be common type I viral fusion proteins TMD topologies (Benton et al., 2018). The arrangement of helices shown here for HIV Env is different than the three-helix bundle topology observed in NMR structures of the TM helices alone (Chen and Chou, 2017; Chiliveri et al., 2018; Dev et al., 2016; Hollingsworth et al., 2018; Kwon et al., 2018a). Thus, the compact three-helix bundle conformation likely represents the low-energy post-fusion conformation of the TMD and its formation is preferred in the minimal constructs used in the NMR and MD studies. More recently, a similar study of isolated MPER-TMD peptide was done in nanodiscs supporting our conclusions that in more native lipid environment stable, trimeric topology is unfavorable (Wang et al., 2019) Our data demonstrate how the ectodomain and MPER restrain the TMD in a crossed topology, particularly apparent in the MPER Fab -bound state. This topology of the TMD is consistent with the meta-stable prefusion state primed for the energetically downhill conformational changes associated with post-fusion conformation. In the nanodisc environment, an additional vertical TMD orientation was observed (Figure 4G). Strikingly similar long, continuous and vertical TMD helix was recently suggested based on molecular dynamics simulation of LN01 antibody in complex with complete TMD (Pinto et. al. 2019). In nanodisc structure presented here, the R696 TMD crossing-point is disconnected from other protomers, suggesting that the TMD domain coordination is influenced by the MPER antibody binding. As compared to micelle, the nanodisc bilayer introduces additional support for the Fab binding, which may result in a more native environment and allow the separation of TMDs in contrast to micelle-embedded complex. The inference is also supported by the difference in position of 10E8 Fab in the AMC011FL nanodisc as compared to the JRFL∆CT micelle, where the membrane-surface-facing side of the docked Fab-MPER peptide structure (PDB: 5T80) is ~10Å closer to the adjacent protomer as compared to the nanodisc (Figure 4I). Therefore, the micelle appears to allow closer juxtaposition of MPER peptides from adjacent protomers when bound to the Fab. Alternatively, the difference in the MPER Fab bound protomer TMD topology could be due to differential effects imposed by the VRC42.01 and 10E8 antibodies.

Our data demonstrate that the MPER is difficult to access and, thus, support a progressive binding model for MPER antibodies that occur in series of steps (Figure 6). Initial contacts of antibody between the Env ectodomain and the lipid surface lead to tilting of Env. This interaction in turn increases the exposure of the MPER peptide by partially lifting it off from the membrane surface (Figure 4D, S4 and S5), lending support that the membrane-embedded and exposed or lifted conformation of the MPER peptide are both relevant for binding (Fu et al., 2018; Irimia et al., 2017; Kwon et al., 2018a). Binding of additional MPER antibodies eventually raises the whole Env up off the membrane and may contribute to shedding of gp120 as shown in earlier studies (Ruprecht et al., 2011). This model would also be in agreement with studies showing increased neutralization efficiency after binding of the CD4 receptor (Kim et al., 2014; Rathinakumar et al., 2012), which results in a steeper angle of HR2, the helix immediately N-terminal to the MPER (Ozorowski et al., 2017). In the context of the Env clustering on the mature virus particle surface and formation of the entry claw (Carravilla et al., 2019; Sougrat et al., 2007), the tilting of two Envs away from each other would also inhibit the formation of the entry claw and possibly contribute to the neutralization efficiency. The tilting component could be mechanistically related to the binding of 35O22, another bNAb that targets an epitope at the interface of gp41 and gp120 and similarly binds better after CD4 receptor engagement (Huang et al., 2014). Finally, given the geometry of the three MPERs in trimeric Env, the two Fab arms of an intact IgG would preferentially bind to two different Env trimers, rather than to a single Env spike as seen on PC64FL bicelle surface with DH511 IgG (Figure S5). Given the limitations of the lipid assembly system, the stoichiometry of MPER antibody arms may however be different in the biological context where membrane surface is circular and Env is more prone to shed of the gp120. Second and third antibody arm binding to same trimer on the virus particle may therefore encounter an Env that has already shed of the gp120 and does not need tilting for accessing the MPER.

**Figure 6.**
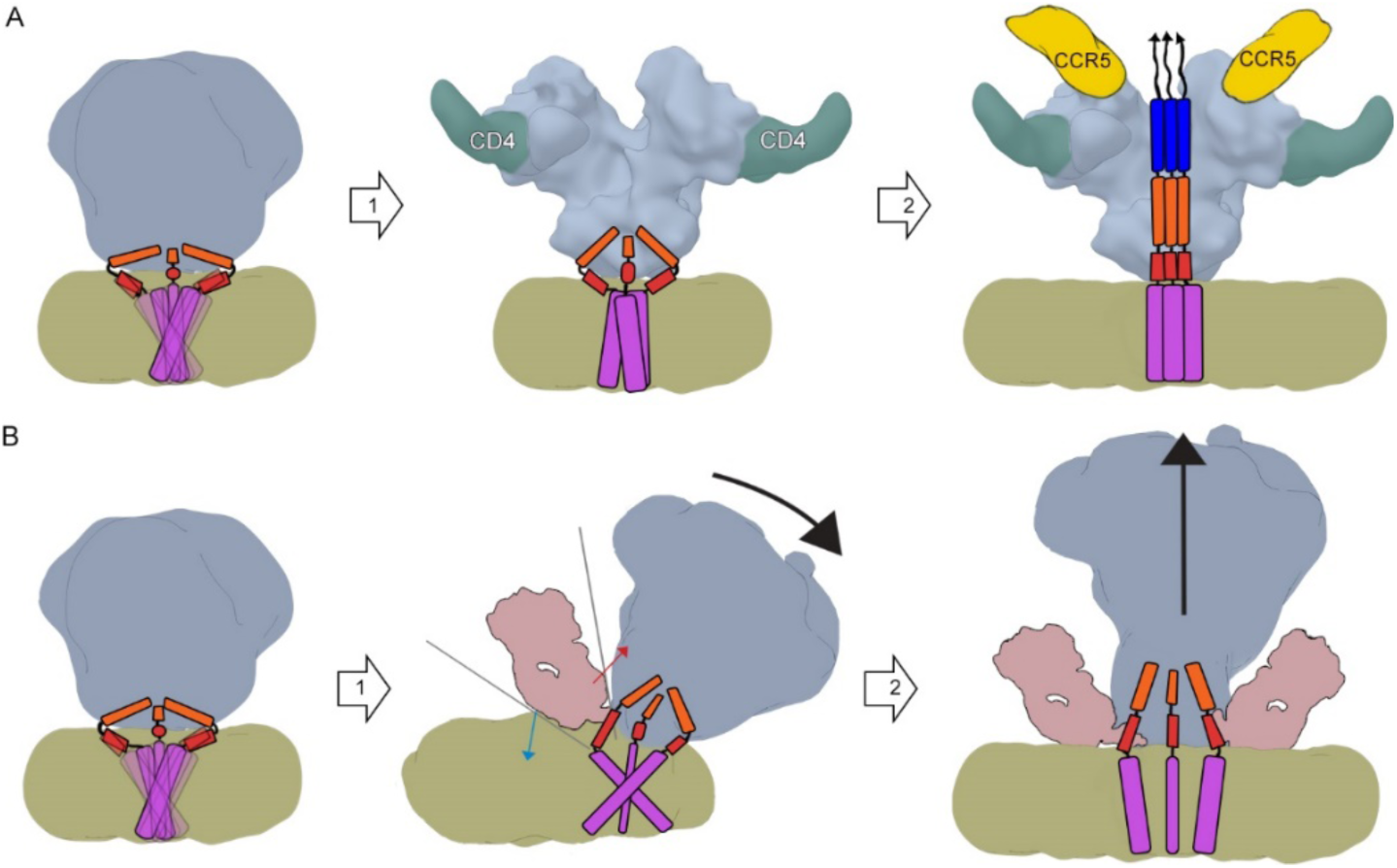
Model for MPER-targeting neutralization mechanism steps based on lipid bilayer systems. Suggested HR1 (blue) HR2 (orange), MPER (red), and TMD (purple) orientations are presented. Based on the lack of density corresponding to the TMD in our EM reconstructions we hypothesize that in their ground state, TMDs are in a loosely folded scissoring fluctuation. A) In the absence of an MPER-targeting Fab, (1) CD4 triggers stabilization and conformational changes in HR2, MPER and TMD in addition to opening of the ectodomain and exposure of the CCR5 co-receptor binding site. (2) This leads to co-receptor binding and formation of a lower energy state post-fusion conformation of fusion peptide (arrows), HR1, HR2, MPER and TMD. Further downstream (not represented in this schematic), this extended three-helix bundle undergoes a further condensation into a six-helix bundle and, in concert with adjacent Env molecules, facilitate membrane fusion and viral entry into the host cell. (B) In the MPER-targeting neutralization path (1), approaching antibody forms a wedge between the ectodomain (red arrow) and bilayer surface (blue arrow), tilting the ectodomain, increasing the exposure of MPER peptide and stabilizing the scissoring of MPER-TMD. While the tilting may restrict access to additional MPER epitopes within the trimer we observe at least a second MPER binding event. (2) Thus, subsequent Fab arms binding to other protomers eventually lead to lifting of Env and possibly increased gp120 shedding in the absence of stabilizing mutations or PGT151 Fab. MPER-TMDs are now separated and locked by the MPER antibody.

The structures with 10E8 yield insight into its mode of binding and possible developmental pathways. As observed in the complex of JRFL∆CT with 10E8 (Lee et al., 2016), our AMC011FL nanodisc complex illustrates that glycans N88 and N625 are positioned to clash with the antibody. These clashes would occur in the proximity of the framework 3 region (FR3) of the 10E8 heavy chain. FR3 likely also has peptide contacts with gp120 and gp41 (table 1) and is a component of 10E8 paratope on the lipid surface and the ectodomain. In another study, 10E8 still showed substantial, albeit reduced, neutralization activity when the FR regions were reverted closer to the germline sequence (Georgiev et al., 2014). Importantly, however, these germline revertants had mature CDRs grafted into the constructs and, therefore, the significance of the FR regions to early antibody maturation events may have been lost. It is also important to note that framework mutations are shown to be generally required for broad neutralization (Klein et al., 2013). Thus, in the light of these studies, our results suggest that immunogens that can drive mutations in FR3 may be important in triggering maturation of MPER bNAbs.

Most MPER immunogen design efforts have focused on using the MPER peptide in isolation (Liu et al., 2018) largely because of the difficulty in producing FL Env. The critical challenges in using transmembrane Env versions are the low protein production level and the instability of the purified transmembrane protein. We achieved ~100-300µg total yield per 1L of 293F cells for stabilized BG505-ST-710, which is roughly an order of magnitude away from what would be required for immunization trials in large animal models. Furthermore, compositional heterogeneity could complicate immunogen formulation. Nonetheless with the BG505-ST-710 peptidisc, we were able to display full MPER epitopes in a single, stable molecule. Stabilizing mutations also allowed the purification protocol to follow closely the methods standardized for soluble Env. Env-nanodisc immunization could then address three important open questions: first, the transmembrane version of Env has been shown to possess a glycan shield that is closer to the composition of the glycan shield presented on virus particles (Cao et al., 2018; Rantalainen et al., 2018; Seabright et al., 2019; Torrents de la Pena et al., 2019). Second, a recent analysis of the polyclonal immune response against soluble SOSIP immunogen indicated that in the non-human primate model, a dominant response targeted the exposed base of the trimer (Bianchi et al., 2018). In membrane-embedded formulations, this immunodominant neoepitope would be protected by the bilayer. Additional prevention of off-target immune responses against the scaffold protein can also be minimized with less immunoreactive peptide scaffolds, such as A22 (Kuai et al., 2017; Kuai et al., 2018) and peptidisc (Carlson et al., 2018). The third important factor in Env-nanodisc immunization would be the capacity to present the full set of quaternary epitopes and which more closely resemble the corresponding epitopes presented on native virus particles. Our structures enabled examination of the full MPER epitope in a quaternary context (Figure 4E, table 1). These advances will enable future studies of Env in its native, membrane-embedded environment, contributing to immunogen design for an effective HIV vaccine.

## Acknowledgements

Authors would like to thank Hannah Turner and Bill Anderson for assistance in microscope management and operation, Charles Bowman and JC Ducom for assistance in EM data management and Lauren Holden for critical reading of the manuscript. DH511.2 Fab and IgG were generous gifts from Barton F. Haynes, Duke University School of Medicine, Durham, NC.

This work was supported by the NIH National Institute of Allergy and Infectious Diseases grants UM1AI100663 and UM1 AI144462: R01 AI143563 (MBZ); by the Bill and Melinda Gates Foundation through the Collaboration for AIDS Vaccine Discovery (CAVD), grant OPP1115782. This is manuscript number 29908.

## Accession numbers

Negative stain and cryo-EM reconstructions have been submitted to Electron Microscopy Data Bank with following access codes. PC64FL+PGT151 Fab+VRC42.01 Fab detergent-lipid micelle (xxxx), AMC011FL+PGT151 Fab+VRC42.01 Fab detergent-lipid micelle (xxxx), PC64FL+PGT151 Fab+VRC42.N1 PC64FL + PGT151 Fab + DH511 Fab detergent-lipid micelle (xxxx), PC64FL + PGT151 Fab + VRC46.01 Fab detergent-lipid micelle (xxxx), AMC011FL + PGT151 Fab + PGZL1 Fab detergent-lipid micelle (xxxx), AMC011FL + PGT151 Fab + 10E8v4-5R 100cF Fab detergent-lipid micelle (xxxx), BG505delCT + PGT151 nanodisc (xxxx), BG505delCT (Ectodomain) + PGT151 Fab nanodisc (xxxx), PC64FL + PGT151 Fab nanodisc (xxxx), AMC011FL + 2 X PGT151 Fab + 1 X 10E8 Fab nanodisc (xxxx), AMC011FL + 2 X PGT151 Fab + 3 X 10E8 Fab nanodisc (xxxx), AMC011FL + 1 X PGT151 Fab + 2 X 10E8 Fab nanodisc (xxxx), AMC011FL + 1 X PGT151 Fab + 3 X 10E8 Fab nanodisc (xxxx), PC64FL bicelle (xxxx), PC64FL + VRC42.01 Fab bicelle (xxxx), PC64FL + DH511 Fab bicelle (xxxx), PC64FL + PGT151 Fab nanodisc (xxxx), PC64FL + VRC42.N1 Fab bicelle (xxxx), AMC011FL + VRC42.01 Fab nanodisc (xxxx), AMC011FL + 10E8 Fab nanodisc (xxxx), BG505-ST-710 peptidisc (xxxx), BG505-ST-710 + 10E8 Fab peptidisc (xxxx). Atomic coordinates of the hybrid model based on AMC011FL + 1 X PGT151 Fab + 3 X 10E8 Fab nanodisc cryo EM map were submitted to Protein Data Bank with accession code xxxx.

## Author contributions

K.R designed the research and developed the lipid assembly methods. K.R., A.A., W-H.S., and J.L.T prepared Env samples. K.R prepared EM samples. K.R., W-H.S., and Z.B. imaged EM data. K.R. and Z.B. processed EM data. A.A and W-H.S engineered Env constructs. A.A. and K.R. adapted lipid assembly method to immunogen production. T.S. performed BLI experiments. X.Z. performed MS analyses. L.Z., A.I., J.C., K.Z., Y.D.K., W.L., C.A.S., and R.V. provided purified IgG and Fabs. K.R., T.S. and X.Z prepared the Figures. K.R. and A.B.W. wrote the manuscript. S.K., P.D.K., N.A.D-R., I.A.W., M.B.Z., J.R.Y III., W.R.S. and A.B.W. provided the resources and supervised the project.

## Methods

### Recombinant protein expression and purification

The MSP1D1 scaffold protein was expressed and purified according to standard *E. coli* recombinant protein expression methods using plasmid available at addgene (https://www.addgene.org, plasmid #20061). Briefly, BL21(DE3) *E. coli* cells were transfected with plasmid expressing the scaffold and grown to an OD_600_ of ~0.8. Expression was induced with 1mM IPTG for ~5 hours. Cells were harvested and MSP1D1 purified by Ni-NTA affinity purification followed by size exclusion using a Superdex 200 increase 10/300 GL column. The MSP1D1 was used without His-tag cleavage throughout the study. Peptidisc scaffold was purchased from Peptidisc Biotech (https://peptidisc.com/).

Env clones PC64M18C043-FL (PC64FL), AMC011FL and BG505∆CT were purified as described previously (Blattner et al., 2014; Rantalainen et al., 2018; Torrents de la Pena et al., 2019). Briefly, HEK293F cells were transfected with 250 µg of Env DNA per liter of cells and supplemented with 62.5 µg/ml of furin DNA to ensure complete cleavage of Env at cell density of 1.6 milj/ml. Cells were harvested 72 hours post transfection. PGT151 with an engineered TEV site between the Fab and Fc was added on cells prior to lysis with DDM containing buffer. Cleared lysate was mixed with protein A matrix and incubated overnight at +4° C. Next, matrix was washed with 50 mM Tris-HCl (pH 7.4), 300 mM NaCl, 0.1% (w/v) CHAPS, 0.03 mg/mL deoxycholate followed by wash with 50 mM Tris-HCl (pH 7.4), 500 mM NaCl, 0.1% (w/v) DDM, 0.03 mg/mL deoxycholate and finally exchanged to buffer with 50 mM Tris-HCl pH 7.4, 150 mM NaCl, 0.1% (w/v) DDM, 0.03 mg/mL deoxycholate and 2 mM EDTA in gravity flow column. Env was eluted by adding ~200 µg of TEV enzyme per liter of cell culture used and incubated for 4h at room temperature. Sample was then concentrated and purified with size exclusion chromatography using a Superose S6i 10/300 column.

BG505-ST-710 DNA construct, codon optimized for mammalian cell expression, was subcloned into a pcDNA3.4 expression vector. 293F cells were co-transfected with BG505-ST-710 and furin DNA vectors (500 and 250 µg per 1 L of cells, respectively) using PEI (PolySciences, Inc). The cells were harvested by centrifugation (3,000 RCF, 30 min, 4 °C) 48 – 96 hours post-transfection and lysed using 25 mM Tris + 300 mM NaCl + 1% Triton-X 100 buffer (pH 7.4) for 2 hours at 4 °C. Cell lysates were cleared by centrifugation (12,000 RCF, 1 hour, 4°C) and subsequent vacuum-filtration (0.22 µm PES filter, Millipore Sigma). Cleared lysates were run over a Sepharose 4B resin (GE Healthcare) with immobilized PGT145, 2G12 or PGT151 antibodies. BG505-ST-710 was eluted off the column using the elution buffer 25 mM Tris + 3 M MgCl2 + 0.05 % DDM + 0.003 % DOC (pH 7.2). Sample was concentrated, buffer-exchanged to the gel-filtration buffer 25 mM Tris + 300 mM NaCl + 0.05 % DDM + 0.003 % DOC and subjected to SEC (Superose S6i 10/300 column).

### IgG and Fab expression and purification

IgGs and Fabs were expressed following standard protocols as follows. Proteins were expressed in FreeStyle 293F cells (Thermo Fisher - R79007). About 25 mL Opti-MEM (Life Technologies – 31985-070) containing ~750 µg DNA (500 µg heavy chain and 250 µg light chain plasmid) for Fab or ~500 µg DNA (250 µg heavy chain and 250 µg light chain plasmid) for IgG was mixed with 25 mL Opti-MEM containing 2,250 µg polyethylene imine MAX (MW 40,000; Polyscience - 24765-1). After incubation for 20 min at RT, the transfection mix was added to 1L cells at a density of ~1.2×106 cells/ml in FreeStyle 293 Expression Medium (Thermo Fisher - 12338018). The cells were incubated at 37°C and 8% CO2 for 6 days. After harvesting the cells, the supernatant, containing IgG or Fab, was filtered and loaded into a HiTrap protein A column (GE Healthcare 17040301, for IgG) or HiTrap KappaSelect column (GE Healthcare Life Sciences – 17545812, for Fab). The column was washed with phosphate buffered saline and eluted with 0.1 M glycine pH 2.7. The fractions were concentrated, and the buffer was changed to 20 mM sodium acetate pH 5.5. The Fab was loaded into a Mono S column and was eluted with a 0 to 60% linear gradient of 1M sodium chloride in 20 mM sodium acetate pH 5.5 buffer. The Fabs were concentrated and stored in 20 mM sodium acetate pH 5.5 at 4°C.

### Lipid stock preparation

To ensure reproducibility, all lipid stock solutions were prepared as follows: lipids were either first dissolved in chloroform or, when available, in solvent used from pre-dissolved ampules. Solvent was evaporated for ~20 minutes under nitrogen gas flow and slow rotation until lipid was deposited as a thin film on the surface of the ampule. Lipids were next re-solubilized into lipid rehydration buffer containing 50 mM Tris (pH 7.4), 150 mM NaCl and 0.1% DDM, and diluted to a final concentration of 1 mM. After 30 min incubation at room temperature, stock solutions were sonicated using stepped micro tip (3 mm), 20 - 25% efficiency and 50 % time cycle until the solution was clear (5-30 mins). Stocks were then aliquoted and stored in −80C° until use for up to 6 months. The following lipids were used throughout the study: 18:1 (Δ9-Cis) 1,2-dioleoyl-sn-glycero-3-phosphocholine (DOPC), 18:1 1,2-dioleoyl-sn-glycero-3-phospho-L-serine (DOPS), 18:1 (Δ9-Cis) 1,2-dioleoyl-sn-glycero-3-phosphoethanolamine (DOPE), 18:1 PA 1,2-dioleoyl-sn-glycero-3-phosphate (DOPA), 18:1 1,2-dioleoyl-sn-glycero-3-phospho-(1’-myo-inositol-4’,5’-bisphosphate) (PIP2(4,5)), 18:1 (Δ9-Cis) 1,2-dioleoyl-sn-glycero-3-phospho-(1’-rac-glycerol) (DOPG), and cholesteryl hemisuccinate (CHS). The lipid mix for disc assembly was prepared by mixing the thawed lipid stocks to final ratio (see assembly below for details), followed by 3 times freeze-thaw cycle with LN2 and room temperature and vortexing between freeze-thaw cycles to ensure even mixing of lipids in DDM micelles.

### Assembly

An overview of protein purification and assembly workflow is presented in Figure 1A. Protein concentrations were calculated using absorbance at 280nm and corrected with protein specific extinction coefficient factors. Purified Env was concentrated to ~1mg/mL using Amicon concentrators with 100 kDa molecular weight cut off (MWCO) prior to mixing with other components. The lipid mixture stock solution was prepared to a total concentration of 1mM. Typical molar ratio for different lipids in the mixture was 40:40:20 (DOPC:DOPS:CHS). MSP1D1 and peptidisc scaffold were diluted to 2mg/mL stocks. Molar ratios of disc assembly components were screened yielding following standard conditions for the assembly mix: 1:8:240 (Env:scaffold:lipid) for MSP1D1 scaffolded discs and bicelles and 1:120:580 for peptidisc. A typical reaction for MSP1D1 scaffolded assemblies consisted of 25 µL of Env, 25 µL of lipids and 10 µL of MSP1D1 scaffold (tot 60 µL) or 25 µL of Env, 50 µL of lipid mix and 25 µL of peptidisc scaffold (100 µL) at concentrations given above. Assembly mixtures were incubated for~30 min at room temperature prior to assembly initiation by addition of bio-beads. Prior to use, the bio-beads were rinsed with methanol for ~1 min followed by three consecutive washes with water (~10 X vol of the biobeads) and stored at +4C° for up to three days. Approximately 50% vol to total reaction volume of bio-beads was added per assembly reaction. The reaction was incubated for 24-48h at +4°C in a rotating mixer followed by transfer to a new tube with a new batch of bio-beads and additional incubation for 24h to maximize detergent removal. Purification of assembled discs was started by separating aggregated protein by centrifugation for 10 min at 13,000 x rcf at +4°C on a tabletop centrifuge. Discs and bicelles were then subjected to a final polishing purification step either by lentil lectin sepharose or size exclusion chromatography. Lentil lectin purification was initiated by mixing the assembled discs with TBS equilibrated, drained matrix (Lentil Lectin Sepharose 4B, GE Healthcare) in a 1:1 sample-to-matrix ratio. The sample was allowed to bind overnight at +4C°, followed by 3 × washes with ~1.4ml TBS in a 1.5 ml test tube. Discs were eluted by three consecutive 1h incubations with TBS + 1 M methyl α-D-mannopyranoside using an equivalent volume of elution buffer to matrix (e.g. 100 µL elution buffer per 1h elution from 100 µL of drained lentil lectin matrix). Three fractions were pooled and dialyzed against Env-nanodisc buffer (20 mM Tris-HCl, pH 7.4, 40 mM NaCl) in a 20 kDa MWCO slide-a-lyzer dialysis units (Thermo Fisher Scientific) to remove methyl α-D-mannopyranoside and to prepare the sample for concentration by water evaporation (Savant DNA120 SpeedVac, Fisher Scientific). The sample was concentrated 5 to 10 times the original concentration depending on required protein concentration, resulting in the final sample for EM analysis. If size exclusion chromatography was used as the final polishing step, the Superose 6 Increase 10/300 column was equilibrated in Env-nanodisc buffer and used according to the manufacturer’s instructions. Unassembled Env, free scaffold, and lipids were separated, but peaks for different Env occupancies in discs were overlapped too much for separation and were pooled and concentrated as with the lectin purification method. The Env incorporation ratio was measured by calculating the amount of Env added to the assembly reaction versus the concentration in the purified final sample. The stability of disc preparations was assessed by measuring concentration of non-aggregated discs and with EM analysis either after storage in +4°C up to 6 months or by flash freezing in LN2 and storing at −80°C.

### Mass spectrometric identification of lipids

Nanodisc solution (final 0.12 mg/mL, 40 μL) was digested with proteinase K (2:1, w/w) in TBS ammonium acetate buffer at 4°C for 2h or overnight, and diluted with final 40% (v/v) acetonitrile. Lipid LC MS/MS analyses were performed using an EASY-nLC connected with an LTQ-Orbitrap Velos mass spectrometer (Thermo Fisher). For each run, about 3 μL digest in 40% acetonitrile was directly loaded to a C8 analytical column (Phenomenex C8, bead diameter 5 μm, pore size 100Å, column inner diameter 100 μm, length 20 cm), and eluted with high %B gradients at 0.4 μL/min. LC buffer A was 0.1% formic acid/ 5% acetonitrile/H2O, and buffer B was 0.1% formic acid/ 95% acetonitrile/H2O.

MS instrument settings were adapted to those used for peptide analysis when applicable: spray voltage 2.5 kV and capillary temperature 325 °C. CID MS/MS spectra were typically acquired using data-dependent or targeted manner in positive and negative ion modes for precursor m/z range 200-2000. Earlier experiments also acquired MS3 spectra to help confirm lipid identity. Data-dependent MS/MS used a top 10 method, in which one MS scan was followed by MS/MS scans of the top 10 most abundant MS peaks, and were measured by the linear ion trap analyzer using enhanced and normal scan speeds respectively. Ion trap scans had AGC on and used 1 micro scan and max ion injection time of 10 ms; MS/MS used isolation width 2 Th, normalized collision energy 35%, activation Q 0.20 and activation time 20 ms. Dynamic exclusion was applied at 1 Th width for 60 s duration and 2 repeats. The lipid-only vesicle sample formed from the input lipids without protein was used as a reference for lipid identification. Lipid identification was based on fragmentation analysis and comparison of MS and MS/MS spectra with those of the lipid-only vesicle sample and literature.

### Bio-layer interferometry

Antigenic profiles of Env constructs were measured on an Octet Red 96 instrument (ForteBio) at 23°C in Octet buffer (1x HBS-EP+ pH 7.4 (Teknova) supplemented with 1 mg/mL BSA). Streptavidin (SA) Biosensors (ForteBio) were preincubated with 10 µg/mL biotinylated Galanthus Nivalis Lectin (Vector Laboratories) in Octet buffer for at least 10 minutes. BG505-ST-710 Env was captured at 3 µg/mL in Octet buffer for 6 minutes. After a 30 second baseline phase, dilution series of Fabs were passed over the sensors for 180 seconds, followed by a 180 second dissociation phase. Lectin sensors were regenerated using three 5-second incubations with 150 mM phosphoric acid. Data from a reference sensor were subtracted from each curve, Y-axes were aligned to the baseline phase, and the data were fit to a 1:1 Langmuir binding model in the Octet Data Analysis software (version 11.0.2.3, ForteBio). Data were exported, normalized to the maximum signal obtained with PGT121, and plotted in Prism for macOS 8.2.1 (Graphpad Software).

### Electron microscopy sample preparation

Concentrations of final assemblies were estimated using extinction coefficient of full-length HIV Env and verified by negative-stain EM thereby not including the absorbance coming from the scaffold protein. In samples where Env nanodiscs and bicelles were complexed with Fabs for negative-stain EM, three times molar excess of purified Fab fragment per Env trimer was incubated with the Env nanodiscs overnight at +4°C and stained as follows: 3 µL of purified disc preparation at 0.03 – 0.06 mg/mL was applied to a 400 mesh size Cu grid, blotted off with filter paper, and stained with 2% uranyl formate for 60 seconds. Cryo-EM grids were prepared differently depending on sample type: for lipid-detergent micelle 10 µL of Env at 5-7 mg/mL was mixed with 1 µL of 1 mM lipid mix and ~6 times molar excess of Fab, followed by gradual detergent removal and lipid replacement with three consecutive additions of 3-5 bio-beads with 1h incubation on ice between each addition. 3 µL of sample was then applied with 0.5 µL of 0.04% A8-35 amphiphol to either 1.2/1.3 Quantifoil or 1.2/1.3 C-flat grids and flash frozen in liquid ethane using Vitrobot mark IV (Thermo Scientific) without wait time, blot force of 0 and blot time of 6-7 sec. PC64FL and BG505∆CT nanodisc samples were frozen on 1.2/1.3 Quantifoil grids overlaid with in-house made thin carbon film at 0.1 – 0.5 mg/mL using Vitrobot without wait time, blot force of −10 and blot time of 2.5 – 3 sec. AMC011FL nanodisc was prepared with lipid ratio 30:23:20:15:10:1:1 (DOPC:DOPS:CHS:DOPE:DOPA:PIP2(4,5):DOPG) and complexed with 10E8 Fab during the final lentil lectin polishing purification step of the nanodisc preparation. Approximately 10 X molar excess of Fab was added to matrix bound nanodiscs, incubated over night at +4°C, and washed extensively with TBS followed by elution, dialysis and concentrating steps as described above for nanodisc assembly. The sample was then frozen on graphene oxide grids (GO on Quantifoils R1.2/1.3, Cu, 400 mesh, Electron Microscopy Sciences) without wait time, blot force of 0, and blot time of 2.5 – 3 sec at 0.1 – 0.2 mg/mL.

### Electron microscopy imaging and data processing

Negative stain EM data was collected on a Tecnai Spirit microscope operating at 120 keV using Leginon automated image collection software (Potter et al., 1999). The image collection parameters are summarized in table S1. Cryo-EM data were collected on a Titan Krios and Talos Arctica operating at 300 keV or 200 keV, respectively, both equipped with a K2 direct electron detector (Gatan) using Leginon. Data were processed using various workflows depending on sample type as summarized in Figures S2, S3 and S4. In summary, particles from negative-stain EM data were picked from using DogPicker (Voss et al., 2009) and exported to particle stacks using Relion 3.0 (Zivanov et al., 2018). 2D and 3D classification was done using cryoSPARC2 (Punjani et al., 2017). After one or two rounds of 2D classification and selection of particles with bicelle or disc-like features, approximately one reference free ab-initio 3D reconstruction was generated per 10,000 particles Ab-initio models were then refined using homogenous refinement option in cryoSPARC2. Cryo-EM data processing was initiated by frame alignment using MotionCor2 (Zheng et al., 2017), followed by contrast transfer function (CTF) calculation by GCTF (Zhang, 2016), particle picking with DogPicker, and extraction with Relion 3.0, or CTF calculation, particle extraction and 2D classification in cryoSPARC2. For detergent-lipid micelle datasets, particle stacks after 2D classification were exported to Relion 3.0 for subsequent processing steps. Here, initial particle coordinates were set by binning all imported particles by 4 and refining them using a 60 Å low-pass filtered Env ectodomain with two copies of PGT151 Fab as the initial model. This initial refinement was then used as a seed for subsequent one to four rounds of 3D classification before unbinning, final refinement, and postprocessing using Relion 3 standard parameters. Multibody refinement was done using the final, postprocessed 3D refinement as the starting model with Relion 3 multibody refinement function (Nakane et al., 2018). Nanodisc reconstructions were done in cryoSPARC2. Briefly, after one or two rounds of 2D classification, selected particles were classified using negative-stain reconstructions from the same sample as seed models with heterogeneous refinement function. This process was repeated using prior models as seeds when further classification was needed. Final, clean 3D classes were refined with non-uniform refinement and for classes that refined beyond 7Å, they were postprocessed with local resolution estimation followed by local filtering. Angles between the membrane bilayer, Fab and Env were measures by placing first axis on the surface of the bilayer, second axis through the center of density in EM density map and fitted Fab structure, and the third through the 3-fold axis of the fitted Env structure. Height from bilayer was estimated as follows. A centroid was placed in the center of last modelled residues (Asp664) of three Env protomers in Chimera. Then, a bilayer modelled with CHARMM-GUI (Jo et al., 2008) was aligned to the EM density corresponding the nanodisc or bicelle. Distance to from the centroid and closest atom in the modelled bilayer was reported as the distance from bilayer. All EM data were visualized and analyzed using USCF Chimera and ChimeraX (Goddard et al., 2018; Pettersen et al., 2004). Hybrid model of AMC011FL nanodisc in complex with PGT151 Fab and 10E8 Fab was generated and deposited to Protein Data Bank with accession code XXXX. This model was generated by rigid body fitting separate gp120 and gp41 subunits and PGT151 variable domain of AMC011 FL structure (PDB 6OLP), and variable domains of 10E8 Fab with MPER peptide (PDB 5T80).

## Supplemental data

**Supplemental figure 1.**
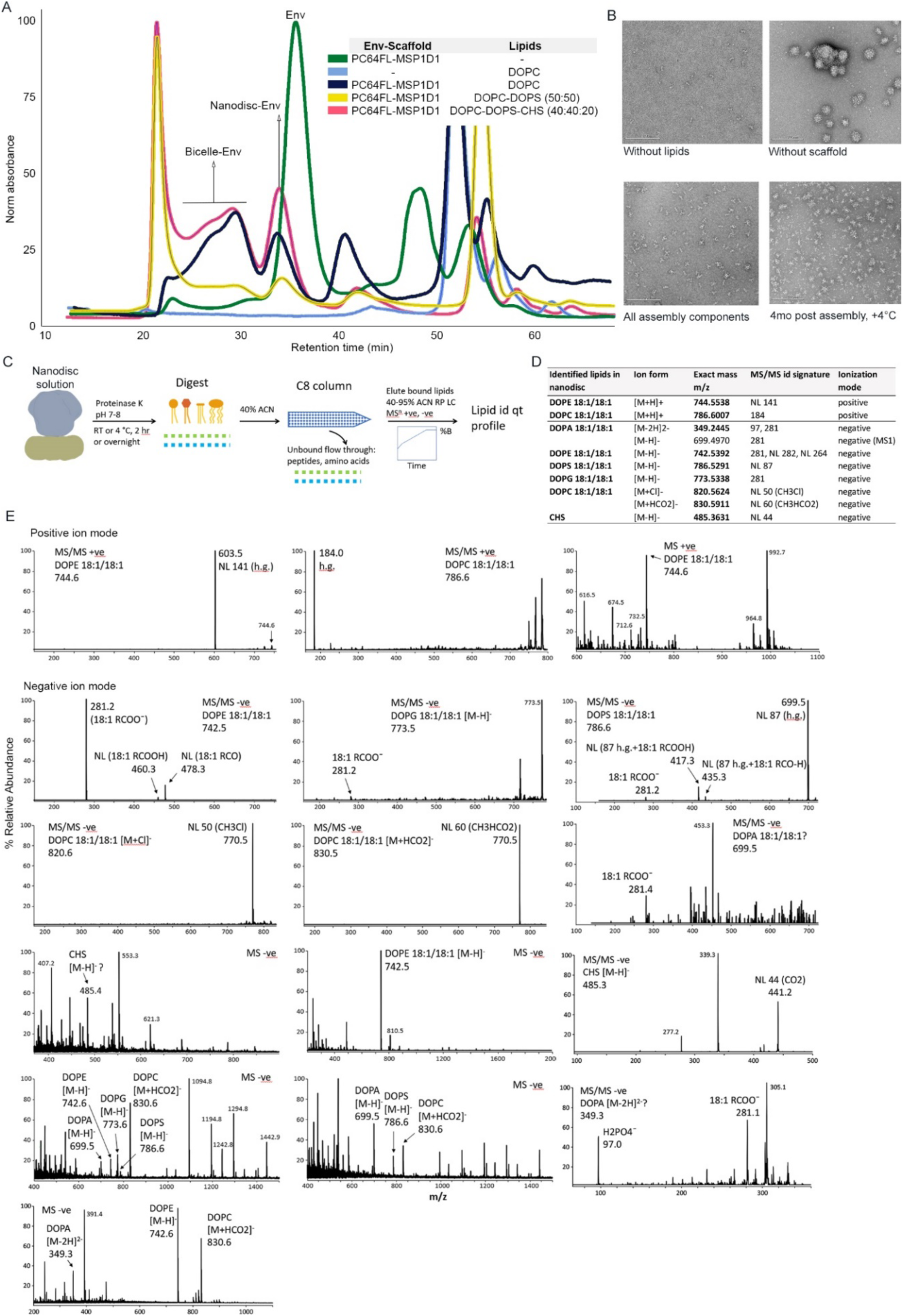
Control of lipid assembly. **Related to figure 1.** A) Representative size exclusion chromatograms from a lipid mixture composition screen. B) representative raw micrographs of control reactions in the absence of scaffold or lipids and when all assembly components were present with an example of discs stored for 4 months at +4°C. C) Digest and conquer lipid analysis workflow. D) Selected disc lipid spectra in positive and negative ion mode. E) Disc lipid identification summary table. NL, neutral loss; h.g., head group; bold font, MS/MS confirmed.

**Supplemental figure 2.**
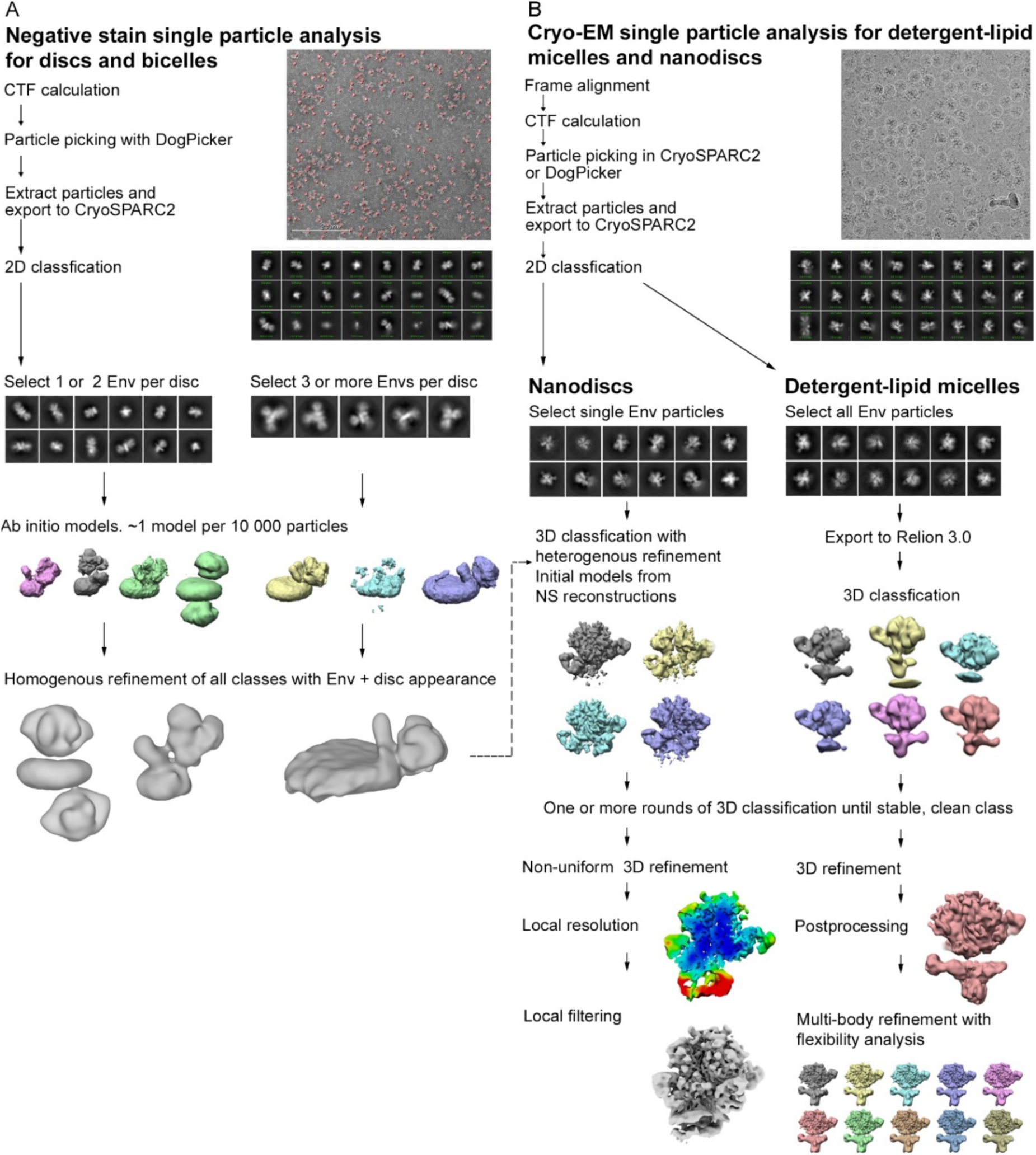
Single particle data processing workflows for analyzing different assemblies. **Related to figures 1, 2, 3 and 4.** A) negative stain EM, and B) cryo-EM.

**Supplemental figure 3.**
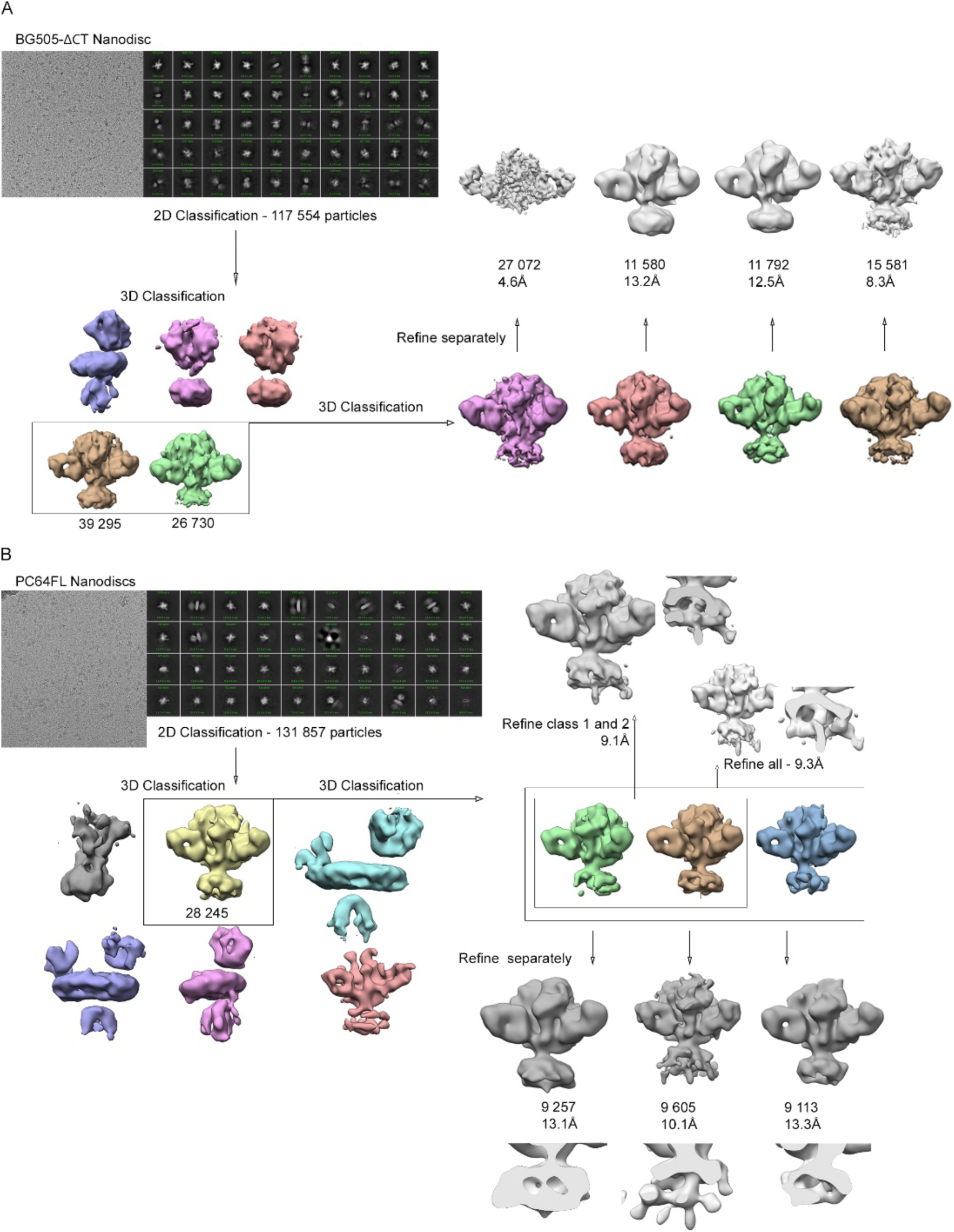
Representative raw micrographs and data processing workflows. **Related to figure 2.** A) BG505∆CT and B) PC64FL nanodiscs. 3D classes with features of PGT151 stabilized ectodomain were selected for further 3D classification and refinement. In PC64FL embedded in nanodiscs, the additional membrane-embedded density was confirmed by refining the subclasses of nanodiscs independently.

**Supplemental figure 4.**
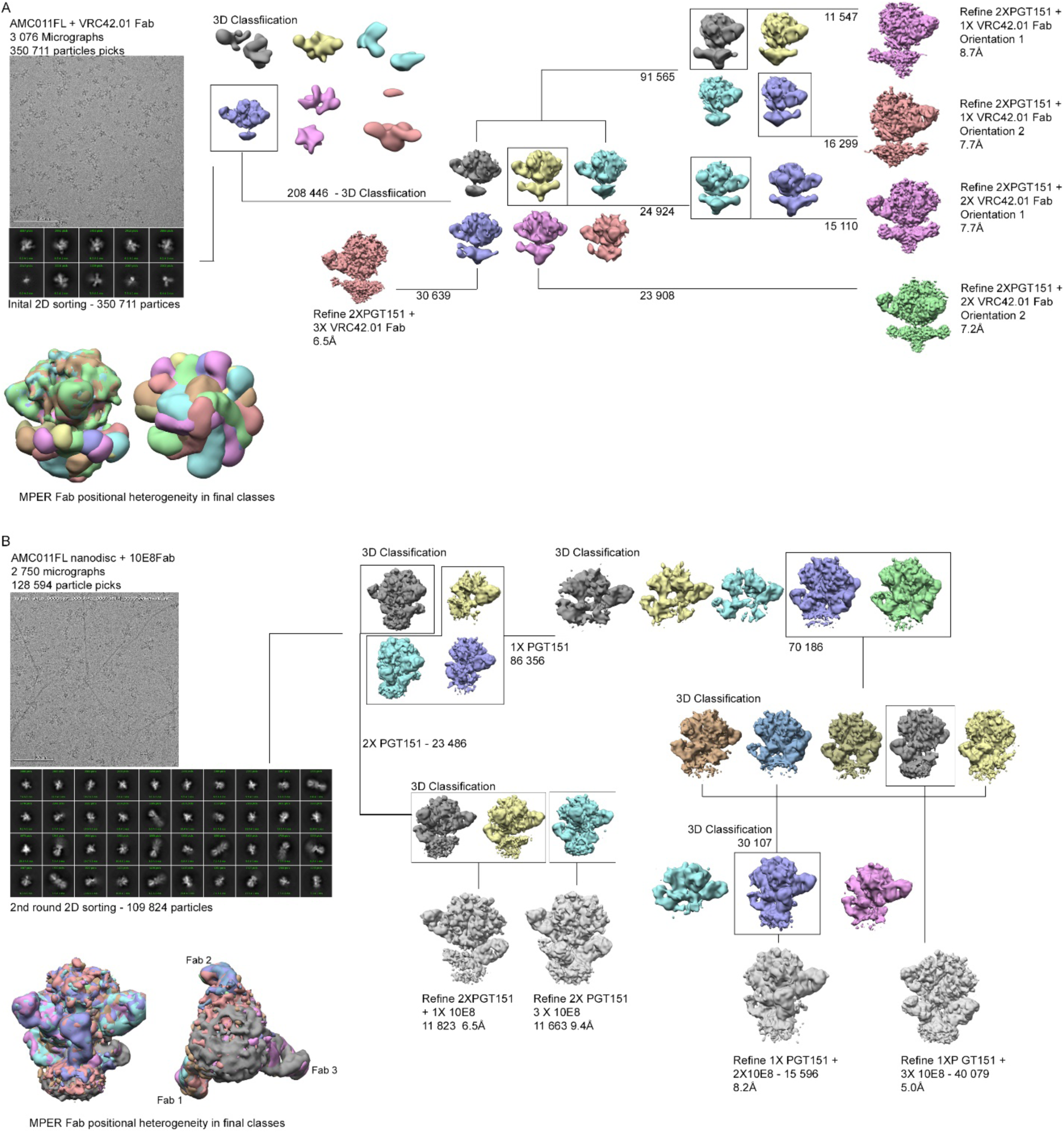
Comparison of data processing workflows. **Related to figures 3 and 4.** A) Detergent-lipid micelle approach and B) nanodisc approach. In the micelle approach, the MPER showed significant positional heterogeneity due to the flexible micelle and leading to near-continuous distribution of Fab positions. In the nanodisc approach, the disc provides additional support for the Fab leading to three distinct and fixed MPER Fab positions. Classification for both approaches were tested in Relion 3.0 and CryoSPARC2. In the micelle samples, different Fab occupancy classes could not be efficiently separated by CryoSPARC2 heterogenous refinement whereas, with the nanodisc approach, both software provided comparable Fab occupancy separation. C) Comparison of PC64FL nanodisc without MPER Fabs and AMC011FL nanodisc with 10E8 Fab with height difference from membrane indicated.

**Supplemental figure 5.**
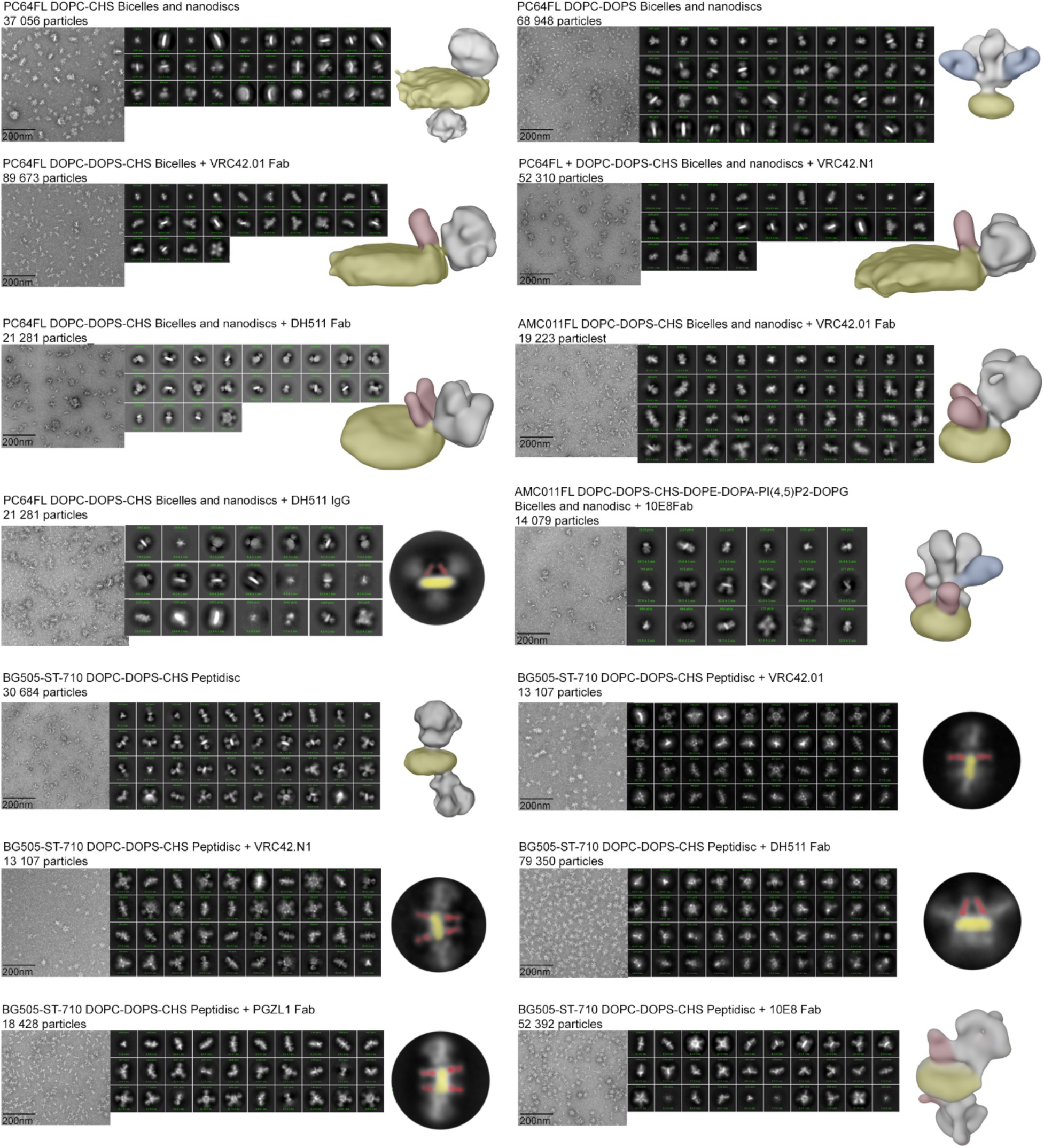
**Related to figures 1, 2 and 4.** A panel of additional complexes were analyzed as controls and as part of the method development. Where data processing did not yield 3D reconstructions, matching features observed in 2D classes and representative 2D classes are shown. PGT151 Fab is highlighted in blue, MPER targeting Fab in red and the lipid bilayer in yellow.

**Supplemental table 1.**
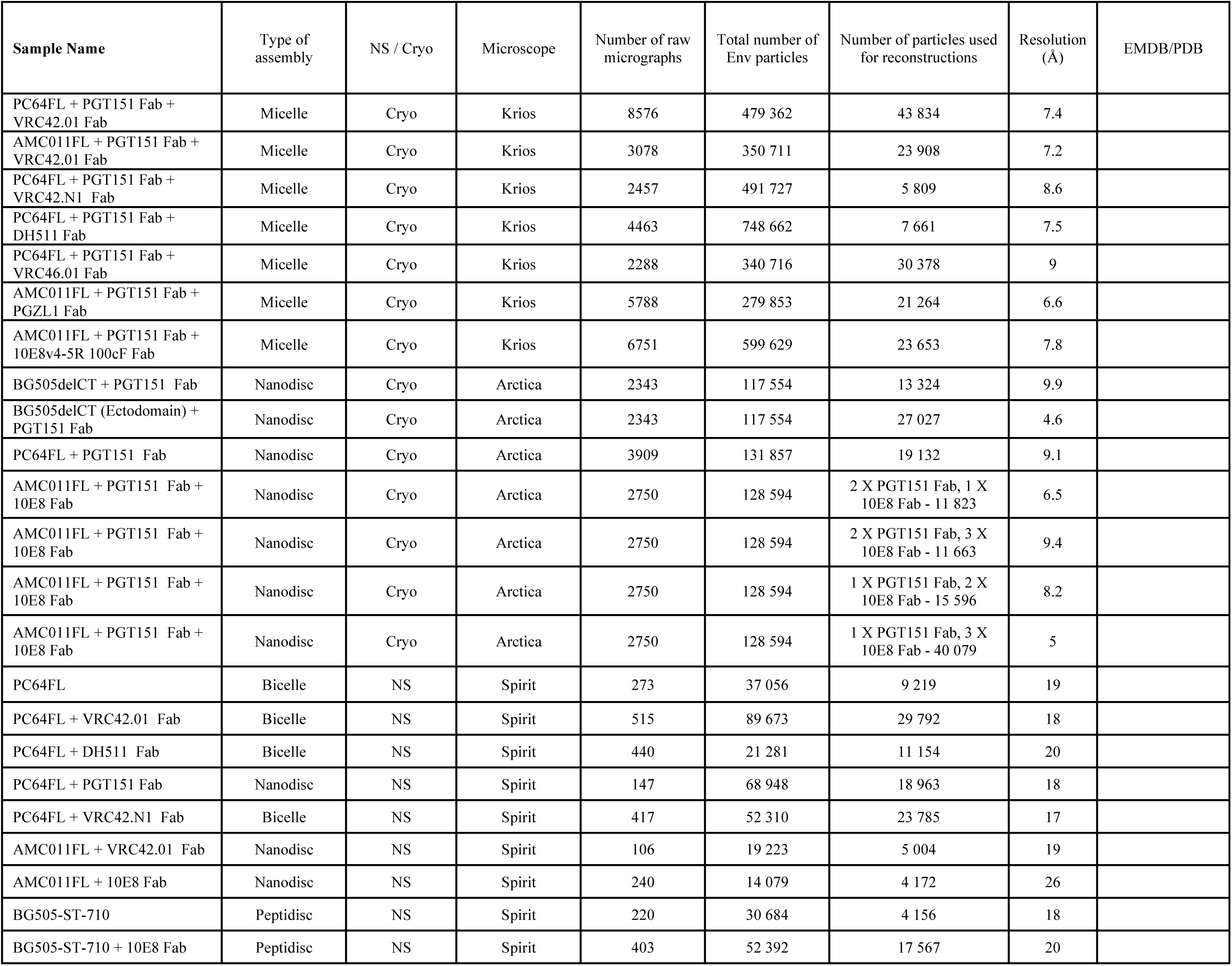
**Related to figures 3, 4 and 5.** Summary of EM samples and imaging conditions.

**Supplemental video 1. Multibody refinement of AMC011FL detergent-lipid micelle in complex with PGT151 Fab and PGZL1 Fab. Related to figures 2 and 3.** Final refinement was used for generating a multibody refinement and principal component analysis of the relative orientations of Env and Fab using Relion 3.0 software. Ten models were generated and combined in sequence in Chimera to illustrate the movement of micelle bound Fab in relation to the Env ectodomain.

